# Estimating consensus proteomes and metabolic functions from taxonomic affiliations

**DOI:** 10.1101/2022.03.16.484574

**Authors:** Arnaud Belcour, Pauline Hamon-Giraud, Alice Mataigne, Baptiste Ruiz, Yann Le Cunff, Jeanne Got, Lorraine Awhangbo, Mégane Lebreton, Clémence Frioux, Simon Dittami, Patrick Dabert, Anne Siegel, Samuel Blanquart

**Affiliations:** Univ Rennes, Inria, CNRS, IRISA, F-35000 Rennes, France; Univ. Grenoble Alpes, Inria, 38000 Grenoble, France; Université Grenoble Alpes, CNRS, LIPhy, Grenoble, France; INRAE, UR1466 OPAALE, 17 Avenue de Cucillé, 35044 Rennes, France; Inria, INRAE, Université de Bordeaux, 33400 Talence, France; Sorbonne University, CNRS, Integrative Biology of Marine Models (LBI2M, UMR 8227), Station Biologique de Roscoff (SBR), 29680 Roscoff, France

## Abstract

**Purpose:** Metabarcoding, and metagenomic sequencing have enabled the characterization of highly diverse environmental communities. The challenge of estimating the metabolic functions carried out by these communities has led to the development of several state-of-the-art methods, most of which are tailored to a specific gene marker. However, the increasing diversity of approaches resulting from advances in sequencing technologies drives the need for methods capable of handling heterogeneous microbial community data. Moreover, predictions often depend on their internal analysis pipelines and are influenced by the underlying databases, which link marker genes to specific functional annotations. This limits users’ ability to evaluate the quality of predictions by tracing internal data and processes. Finally, users are constrained by the specific annotations provided by these methods (e.g. EC numbers), limiting their ability to conduct further specialized analyses based on intermediate results.

**Methods:** EsMeCaTa predicts consensus proteomes and their associated functions from taxonomic affiliations. A key feature of EsMeCaTa is its explainability and flexibility. To support the flexible integration of heterogeneous sequencing data, EsMeCaTa utilizes taxonomic affiliations obtained through analyses of diverse sequencing datasets. To provide insight into the knowledge available for each taxonomic affiliation and to interpret the relevance of predicted functions, EsMeCaTa identifies a taxonomic rank within a given affliation that is suffciently represented by documented proteomes in the UniProt database. The proteins of the UniProt proteomes are clustered and filtered according to a threshold to create consensus proteomes. These consensus proteomes are automatically annotated with functional information (e.g., EC numbers, GO terms) but they are also designed to be used in further customized annotation workflows. Functional annotations are reported in a functional table, which can be enriched with taxon abundances to generate comprehensive functional profiles.

**Results:** EsMeCaTa predictions have been validated using multiple datasets and compared to a state-of-the-art method. Additionally, it was applied to a novel metabarcoding dataset from a methanogenic reactor, characterizing the microbial community and biogas production across different time points and intake condition. Our results demonstrate the link between biogas production, intake condition and the dynamics of the metabolic functions predicted by EsMeCaTa in the microbial communities.

## Background

Metabarcoding and metagenomic sequencing have allowed the characterisation of environmental communities, such as human [1], soil [2] or marine [3] microbiota. While metabarcoding focuses on the sequencing of an amplified marker gene of interest (also named amplicon), metagenomic sequencing provides broader information about the entire genomic content of the sample, allowing for the assembly and binning of genomes [4]. The growth of such sequencing data has led to the creation of open access databases, such as MGnify [5, 6], which provides a unique overview of the availability of environmental sequencing data. For example, 480,962 amplicon data, 57,629 assemblies and 39,920 metagenomes are available from MGnify in 2024^1^. Estimating the metabolic functions performed by the community, its functional profile, is an important issue. HUMAnN3 [7] creates functional profiles directly from the metagenomic sequencing data. For amplicon data, too, several methods have been developed to create functional profiles (called in this article *functional profiling methods*): PICRUSt/PICRUSt2 [8, 9], Paprica [10], Tax4Fun/Tax4Fun2 [11, 12], Piphillin [13, 14], MicFunPred [15] or PanFP [16]. In these cases, a preliminary task is to estimate taxonomic affiliations of the amplicons. Then functions are associated with the taxa or with the sequences directly. Finally, the functions are scaled by the abundances of the taxa in the sample to produce the functional profile.

Functional profiling methods rely on marker gene to predict the functions. One of the first step is to place the gene marker sequences inside a reference space associated with genomic data to find the closest related organisms. The 16S rRNA gene is one of the earliest marker genes used to analyse the bacterial diversity in environmental communities [17] and is currently the most widely used marker gene. Therefore, several methods (such as PICRUSt2) focus their input on this gene and compute functional profiles according to a curated internal database. However, other genes have shown interesting performance in taxonomic characterisation, such as the *rpob* gene [18]. In addition, there are other sequencing methods that provide taxonomic characterisation from environmental samples, such as shallow whole genome sequencing [19, 20] or metatranscriptomics [21]. In this context, an appropriate strategy to deal with the heterogeneity of these sequencing data is to use their common output feature: the *taxonomic affiliations* of the community, *i.e.* all taxa from the taxonomic lineage (from highest taxonomic rank, such as superkingdom, to lowest taxonomic rank, such as genus) of each identified organism, as done in PanFP [16].

The estimation of functions associated with a given taxon relies on comparative genomics approaches applied to the available related reference genomes. PICRUSt2 uses genomes from the IMG database [22], Paprica from the NCBI Genbank database [23] and PanFP from the NCBI Genomes resource [24]. The predictions for a given taxon are strongly influenced by the available information associated with the known organisms and the phylogenetic distances between the identified organisms and the closest reference genomes. For metabolic annotation, the greater this phylogenetic distance is, the lower the accuracy of the predicted functions is [25]. This favors methods that allow users to filter relevant predictions, *e.g.* according to the distance to the closest organisms with available genomes.

The availability of reference genomes is not the only factor impeding the prediction accuracy. Indeed, functional profiling methods rely on an annotation database to link the selected genomes with specific functions and generate functional annotations with internal tools [8, 9, 16]. A first issue with respect to these local databases is that functions are predicted from the genomes during the creation of the database, implying that their predictions may not be up-to-date with respect to current knowledge. A second issue is that the types of predicted functions are limited to the ones selected by the method (for example, EC numbers, KEGG Orthologs and MetaCyc pathways), preventing users to enrich annotations with their own dedicated annotations tools. These issues prompted us to develop a method complementary to currently available methods, providing to the users more flexibility for function prediction by giving access to the sequences leading to the prediction of the taxa functions.

This paper presents EsMeCaTa, a method which predicts consensus proteomes and their associated functions. To allow a flexible management of heterogeneous sequencing data, EsMeCaTa uses as input taxonomic affiliations preliminary obtained through either barcoding, metabarcoding or metagenomics sequencing data. To give insight into the available knowledge for each taxonomic affiliation and interpret the relevance of the predicted functions, EsMeCaTa selects a taxonomic rank in the affiliation of a taxon for which enough proteomes are documented in the UniProt database. As a key output towards explainability and user flexibility, EsMeCaTa creates *consensus proteomes*, the consensus sequences created from the clustering of UniProt proteomes at a specific taxonomic rank. These sequences are automatically associated with functional annotations (EC numbers, GO terms, Kegg IDs) but they also aim to be integrated into further customized annotation approaches. Functional annotations are reported in a *function table* which can be further enriched with taxon abundances (when available) to create functional profiles. Altogether, EsMeCaTa ensures explainability by comprehensively reporting information at each step of the pipeline, such as taxa metadata, Uniprot proteomes, UniProt protein IDs, consensus sequences and eggNOG-mapper predicted annotations.

EsMeCaTa was benchmarked using an algal microbiota dataset comprising paired 16S rRNA sequences and genomes, along with four large-scale microbial community datasets retrieved from the MGnify database. These datasets encompassed diverse environments, including marine microbiota and host-associated microbiota. The benchmarking assessed the accuracy of EsMeCaTa against metagenomes in predicting functions from taxonomic affiliations and evaluated the relevance of its consensus proteome predictions. EsMeCaTa was also compared to the sate-of-the-art method PICRUSt2. The results demonstrated the robustness, accuracy and flexibility of EsMeCaTa, particularly at the species, genus, and family taxonomic ranks. To further showcase its utility, we applied EsMeCaTa to a case study involving a novel dataset sampled from a methanogenic reactor at different time points. This application enabled the identification of functions specific to distinct taxonomic groups by conducting enrichment analyses on EsMeCaTa predicted annotations. It allowed the exploration of three methanogenic pathways based on the predicted functions and consensus proteomes leading to the classification of OTUs based on their enzymatic potential to perform these pathways. This analysis demonstrated that biogas production measurements could be explained by the combination of different methanogenic pathways, performed by different archaeal taxa.

## Results

### EsMeCaTa: Predicting organism functions from taxonomic affiliations

Using the organisms’ taxonomy, publicly-available proteomes and comparative genomics, EsMeCaTa provides estimations of metabolic capacity from taxonomic affiliations.

### Method overview

We first illustrate the method on a set of 13 different taxa selected to cover both prokaryotic (*Gammaproteobacteria*) and eukaryotic (*Alveolata*) taxa of diverse taxonomic ranks (from clade to genus). These examples illustrate the amount of available proteomes and the biases toward most studied groups (column "Input taxa" in Table 1, see Methods).

**Table 1.**
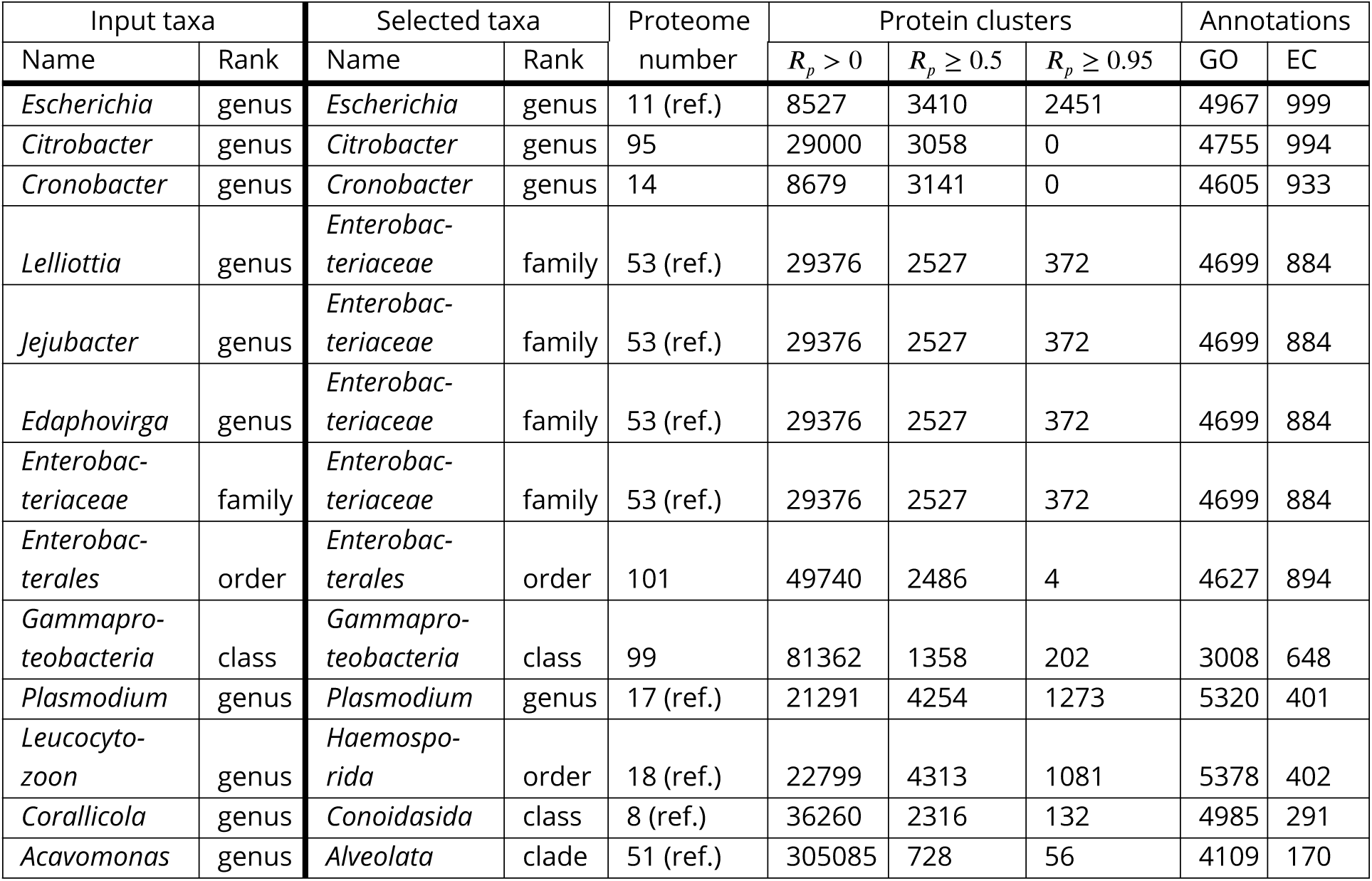
Predictions of EsMeCaTa on the *toy example dataset*. In column "Proteome number", "ref." denotes the use of reference proteomes only, in the case there was at least 5 reference proteomes, otherwise the reference proteomes were used with other proteomes. In column "Protein clusters", 𝑅_𝑝_ frequency of the cluster’s proteins among proteomes, when superior to 0, it corresponds to all protein clusters found by MMseqs2 (thus similar to a pan-genome), at value 0.5, it corresponds to the default threshold used by EsMeCaTa and when superior or equal to 0.95, it corresponds to the notion of relaxed core-genome (protein clusters found in almost all proteomes).

| Input taxa |  | Selected taxa |  | Proteome number | Protein clusters |  |  | Annotations |  |
| --- | --- | --- | --- | --- | --- | --- | --- | --- | --- |
| Name | Rank | Name | Rank | | $R_p > 0$ | $R_p \geq 0.5$ | $R_p \geq 0.95$ | GO | EC |
| *Escherichia* | genus | *Escherichia* | genus | 11 (ref.) | 8527 | 3410 | 2451 | 4967 | 999 |
| *Citrobacter* | genus | *Citrobacter* | genus | 95 | 29000 | 3058 | 0 | 4755 | 994 |
| *Cronobacter* | genus | *Cronobacter* | genus | 14 | 8679 | 3141 | 0 | 4605 | 933 |
| *Lelliottia* | genus | *Enterobacteriaceae* | family | 53 (ref.) | 29376 | 2527 | 372 | 4699 | 884 |
| *Jejubacter* | genus | *Enterobacteriaceae* | family | 53 (ref.) | 29376 | 2527 | 372 | 4699 | 884 |
| *Edaphovirga* | genus | *Enterobacteriaceae* | family | 53 (ref.) | 29376 | 2527 | 372 | 4699 | 884 |
| *Enterobacteriaceae* | family | *Enterobacteriaceae* | family | 53 (ref.) | 29376 | 2527 | 372 | 4699 | 884 |
| *Enterobacterales* | order | *Enterobacterales* | order | 101 | 49740 | 2486 | 4 | 4627 | 894 |
| *Gammaproteobacteria* | class | *Gammaproteobacteria* | class | 99 | 81362 | 1358 | 202 | 3008 | 648 |
| *Plasmodium* | genus | *Plasmodium* | genus | 17 (ref.) | 21291 | 4254 | 1273 | 5320 | 401 |
| *Leucocytozoon* | genus | *Haemosporida* | order | 18 (ref.) | 22799 | 4313 | 1081 | 5378 | 402 |
| *Corallicola* | genus | *Conoidasida* | class | 8 (ref.) | 36260 | 2316 | 132 | 4985 | 291 |
| *Acavomonas* | genus | *Alveolata* | clade | 51 (ref.) | 305085 | 728 | 56 | 4109 | 170 |

The method takes as input a tabulated text file containing taxonomic affiliations compatible with the NCBI Taxonomy [26], that is, the taxonomic lineage describing an organism going from the highest taxonomic rank (such as the Kingdom) to the lowest one possible (such as the species or genus). The input can be the result of different analyses such as taxonomic assignments of marker genes (such as 16S rRNA gene), manually selected taxa or taxa identified from genomes or metagenomes.

The method is divided into three steps (described in following paragraphs and in Fig 1), each yielding a global survey of the current knowledge about each taxon of interest. First, the pipeline retrieves on UniProt [27] the proteomes associated with each taxonomic affliation (Fig 1, step 1 "proteomes"). Second, it estimates the clusters of homologous proteins shared by the proteomes using MMseqs2 [28]. EsMeCaTa filters the protein clusters according to a threshold 𝑇_𝑟_ (Fig 1, step

**Figure 1.**
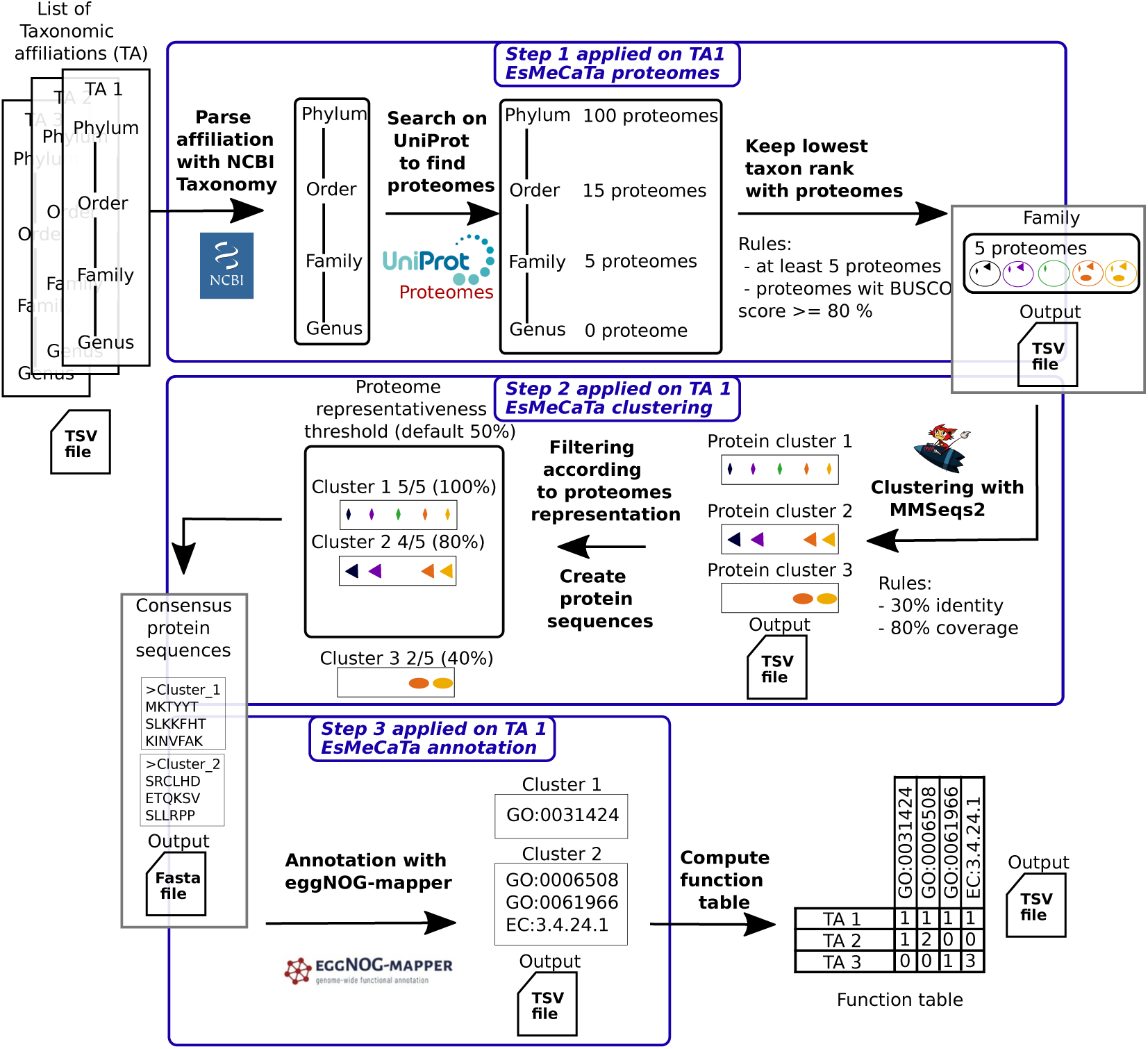
Method workflow. The input to EsMeCaTa consists in a list of taxonomic affiliations (denoted TA in this figure) compatible with the NCBI taxonomy (figure top left). Step 1 is referred to as "EsMeCaTa proteomes" and consists in selecting for each of the input taxonomic affiliations the lowest possible taxonomic rank such that a defined number of proteomes is available in the UniProt proteome database. Step 2 called "EsMeCaTa clustering" consists in computing the clusters of homologous proteins shared across the selected proteomes using MMseqs2, and then in filtering the clusters whose proteins are shared by at least half of the proteomes. Step 3 denoted as "EsMeCaTa annotation" consists in annotating the consensus proteins of each filtered protein clusters using eggNOG-mapper, which provides as output the predicted annotations such as Enzyme Commission (EC) numbers and Gene Ontology (GO) terms (figure bottom right). The predictions for all the different taxonomic affiliations of a dataset are then merged in a *function table* showing the occurrence of the functions according to the predictions made by EsMeCaTa.

2 "clustering") leading to the creation of a *consensus proteome* for each taxon. Third, the consen-sus proteomes are annotated using eggNOG-mapper [29, 30], providing the predicted functions for the taxon (Fig 1, step 3 "annotation"). From the latter predictions, a function table is created summarising the occurrences of annotations (EC numbers and GO Terms) in each taxon. This func- tion table is a key output for the user to undergo further specialized analyses, at both taxon and community scales.

All three EsMeCaTa steps give insights into the number of available related proteomes (columns "Selected taxa" and "Proteome number" in Table 1), the clustering of their proteins (column "Pro- tein clusters" in Table 1), and the functions associated with these protein clusters (column "Anno- tations" in Table 1).

### Step 1: Accounting for the knowledge available about the taxa

For a given taxonomic affiliation (for example, *cellular organisms; Bacteria; Proteobacteria; Gammaproteobacteria; Enterobacterales; Enterobacteriaceae; Escherichia* for genus *Escherichia*), EsMeCaTa searches for the associated proteomes in the UniProt proteomes database [27]. It filters out proteomes with BUSCO score lower than 80% [31]. It selects the taxon associated with at least 𝑁 proteomes and having the lowest taxonomic rank as defined by the input affiliation (𝑁 ≥ 5 proteomes are considered by default). UniProt flags some proteomes as reference proteomes, which are landmarks in the proteome space of organisms. EsMeCaTa first considers reference proteomes and use them if there are at least 5 reference proteomes, otherwise it uses reference and non-reference proteomes. If there are less than 5 reference and non-reference proteomes for the taxon, the method performs again this search with a higher taxonomic rank.

In the example Table 1, six genera in the family *Enterobacteriaceae* are considered. For the genera *Escherichia*, *Citrobacter* and *Cronobacter*, sufficient proteomes are available at the genus level. In contrast for genera *Lelliottia*, *Jejubacter* and *Edaphovirga*, predictions have to be made at the next higher rank, family, to obtain enough proteomes. All three genera are represented by the same proteomes from the *Enterobacteriaceae* family, and hence, are described by the same functions (columns "Selected taxa" in Table 1).

Likewise the four eukaryote genera exemplified as input to the method belong to a heterogeneously studied clade, the *Alveolata*. Within this clade, genus *Plasmodium* is a well-studied organism and gathers 17 reference proteomes, so that predictions can be drawn at this genus level. For the other three illustrated genera (*Leucocytozoon*, *Corallicola* and *Acavomonas*) few or no proteomes are available in UniProt and predictions have to be drawn from higher taxonomic ranks, order *Haemosporidia*, class *Conoidasida* and clade *Alveolata* (columns "Selected taxa" in Table 1). Note that proteomes of the most studied taxa, *e.g.* genera *Escherichia* and *Plasmodium* in the latter examples, are over represented in the predictions made at higher taxonomic ranks. For example, the 18 proteomes associated with order *Haemosporidia* include the 17 proteomes of *Plasmodium*. This also means that certain proteomes can be used several times, for the functional predictions as illustrated in Fig 2A.

**Figure 2.**
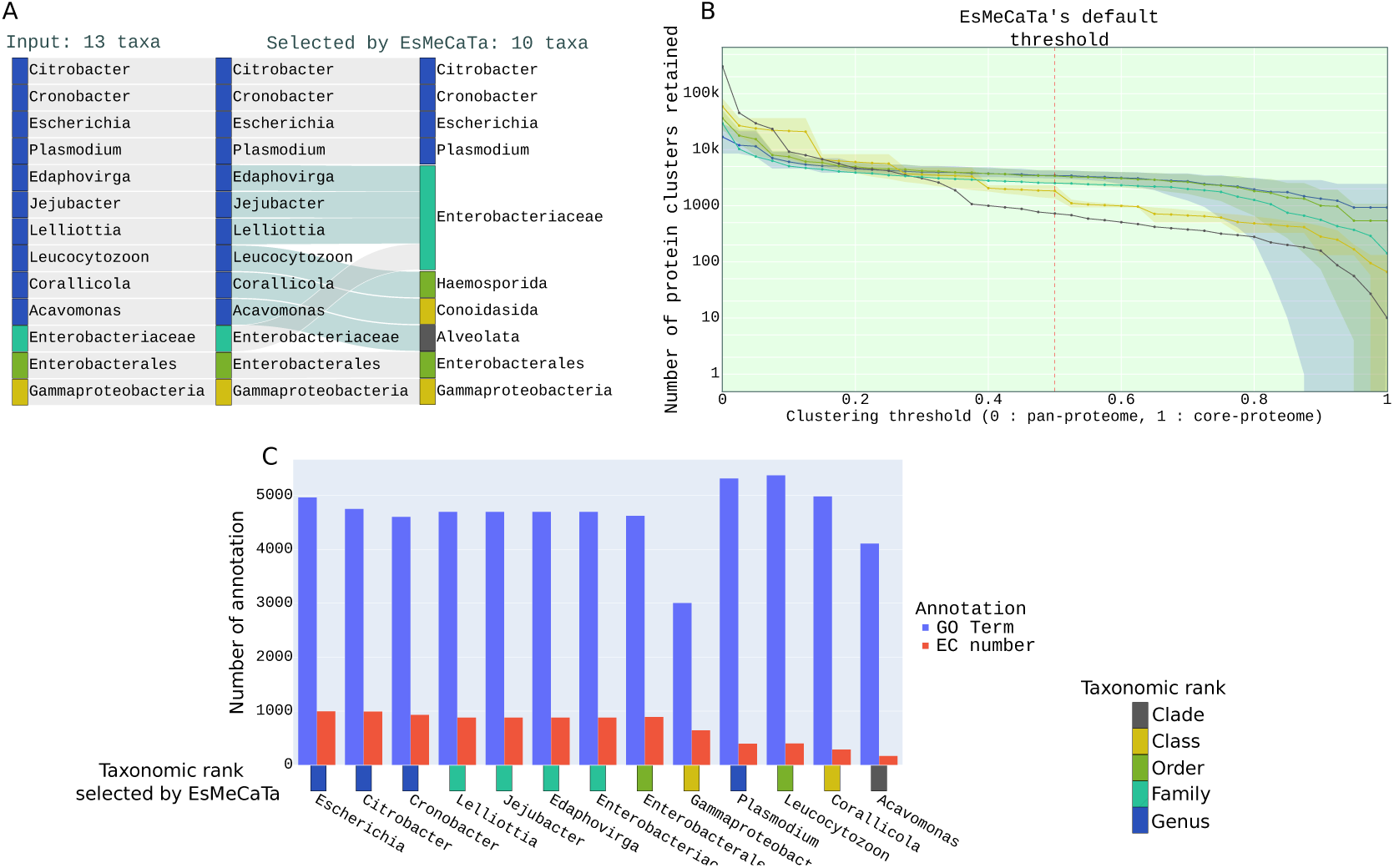
Report on the knowledge available to draw the predicted functions. (A) The Sankey diagram represents the taxa from the *toy example dataset* considered as input (left side) and the corresponding taxa selected according to the proteome availability in the UniProt proteome database (right side). (B) Cumulative distributions of the protein cluster numbers (graph Y-coordinates) according to the proportion of proteomes sharing the clusters (X-coordinates), with the core proteome size indicated at the right (𝑇_𝑟_ = 1) and the pan-proteome size at the left (𝑇_𝑟_ = 0). (C) Number of annotation (EC number and GO Term) inferred for each taxon.

### Step 2: Estimating the protein families shared across the proteomes of a given taxon

The proteins from the proteomes selected at the previous step are grouped into protein clusters using MMseqs2 [28]. Then, the frequency of each protein cluster among proteomes is estimated and denoted as the cluster’s *representativeness* 𝑅_𝑝_. Following the terminology applied in pangenomics [32], the core proteome corresponds to the subset of clusters whose proteins are shared by all the proteomes belonging to a taxon (representativeness 𝑅_𝑝_ = 1). The pan-proteome stands for all the clusters found in at least one proteome, thus having a representativeness 𝑅_𝑝_ > 0.

The distribution of protein clusters over the selected proteomes displayed global trends in line with current findings in pangenomics (columns "Protein clusters" in Table 1, Fig 2B). In particular the relaxed core proteomes (𝑅_𝑝_ ≥ 0.95) consisted in a few thousand protein clusters at the genus level, 2,451 for the genus *Escherichia* and 1,273 for genus *Plasmodium* (Table 1). This was congruent with the core genome sizes reviewed in [32], ranging from 522 to 2,811 in six bacterial genera. The core proteome estimation was sensitive to the quality of the proteomes retrieved in the database, as shown by the two genera *Citrobacter* and *Cronobacter* for which no protein cluster was shared by more than 𝑅_𝑝_ ≥ 95% of the selected proteomes. The method also selected non-reference proteome for these two genera, having BUSCO scores greater than 80%. The proteins potentially missing in several proteomes lead to decreasing the core-proteome size. Along with lesser quality proteomes considered, the higher number of proteomes analysed for genera *Citrobacter* and *Cronobacter* (95 and 14 proteomes respectively, Table 1) would contribute to estimating empty core proteomes. Finally, the obtained core proteomes were smaller when higher taxonomic ranks were considered (Fig 2B), due to the higher taxonomic diversity. This was congruent with previous estimations of the core genomes in class *Bacilli* and phylum *Chlamydiae* involving 143 and 560 genes, respectively [32].

At the genus level, the estimated pan-proteomes included from 8,527 to 29,000 protein clusters (Table 1), consistent with the estimation of pan-genomes ranging from 3,320 to 12,483 gene families in nine bacterial species [33] (containing *Escherichia coli*). Moreover, a wider taxonomic diversity induces a wider gene family diversity: we consistently observed that the higher the considered taxonomic rank, the larger the estimated pan-proteome (Fig 2B).

Finally, EsMeCaTa applies a threshold of 𝑇_𝑟_ = 0.5, meaning that protein clusters are considered for the next annotation step if their proteins are represented in at least half of the selected proteomes (𝑅_𝑝_ ≥ 0.5). This threshold has been chosen based on validations with bacterial proteomes presented below, and can be parameterised by the users. For each protein cluster retained, a consensus sequence is computed. Altogether the consensus sequences constitute the *consensus proteome*.

### Step 3: Predicting the functions of the taxa from the protein clusters

Consensus sequences are then annotated using eggNOG-mapper [29, 30] (see Methods). For each taxon, a tabulated file containing the predictive annotations is created. Finally, these results are summarised into the *function table*, a matrix displaying the occurrence of each annotation (EC number and GO Terms, denoted hereafter as *predicted functions*) in the different taxa (columns "Annotations" in Table 1 and Fig 2 C).

The function table can be examined thanks to several proposed representations. For example, hierarchical diagrams summarise the functions predicted for the taxa, according to the EC numbers (Sup Figure S1). Predictions are also suitable for analyses using dedicated tools, such as function representation using Brenda [34] or Revigo [35], and enrichment analysis using GSEApy [36] and Orsum [37]. In the last sections of the manuscript, we illustrate how the functions predicted from metabarcoding data can be investigated using enrichment analysis and pathway profiling.

### Assessing EsMeCaTa on marine environmental samples and pig, bee and human host-associated microbial communities

Predictions of EsMeCaTa (both consensus proteomes and function tables) were assessed using several microbial datasets from environmental data. EsMeCaTa was applied to the *Ectocarpus sp. microbiota dataset* and the *MGnify dataset* (see Methods).

A first comparison focused on an internal assessment of the cluster filtering threshold 𝑅. This experiment required to launch EsMeCaTa multiple times with different cluster filtering threshold 𝑅, so it was done on a dataset of 10 bacterial complete genomes and 35 MAGs from symbionts [38, 39].

Further comparisons were then performed using the *MGnify datasets* [5, 6] to assess the method accuracy. A comparison with a state-of-art method for predicting functional profiles (PICRUSt2) was performed. A third comparison was made by using all the MAGs with a completeness greater than or equal to 90% to check the quality of predictions made by EsMeCaTa based on three features: (1) EC number (as in previous comparison), (2) Gene Ontology terms and (3) protein sequences. Finally, a fourth comparison assessed the quality of the consensus proteomes predicted by EsMeCaTa according to the corresponding proteins from the MAGs.

### The cluster filtering threshold of 𝑇_𝑟_ = 0.5 as a balance between recall and precision for EC prediction on complete genomes and MAGs dataset

To explore the impact of the cluster filtering threshold on EsMeCaTa prediction accuracy, an experiment was performed on a dataset combining 10 bacterial complete genomes from [38] and 35 MAGS having at least 90% of completeness from [39] (the *Ectocarpus sp.* microbiota dataset, see Method). Five runs of EsMeCaTa were performed on this dataset with five different values of the *representativeness* thresholds 𝑇_𝑟_ (0, 0.25, 0.5, 0.75 and 0.95). The predicted EC numbers for each proteome predicted by EsMeCaTa were compared to the EC numbers from the associated genome annotations. A confusion matrix was then created and F-measures, precision and recall were computed.

Cluster filtering threshold (𝑇_𝑟_ = 0) corresponding to pan-proteome is associated with the lowest precision but the best recall (Figure 3 A). This is expected as, by definition, the pan-proteome contains all the protein clusters of a taxon, and thus a maximum number of true positives and false positives. Conversely, cluster filtering threshold (𝑇_𝑟_ = 0.95) corresponding to core proteome is associated with the highest precision but the lowest recall. There is in this case a limited amount of protein clusters kept, but which are widely represented in the taxon, thus inducing a low false positive rate. The threshold 𝑇_𝑟_ = 0.5 corresponded to a balance between precision and recall (Figure 3 A).

**Figure 3.**
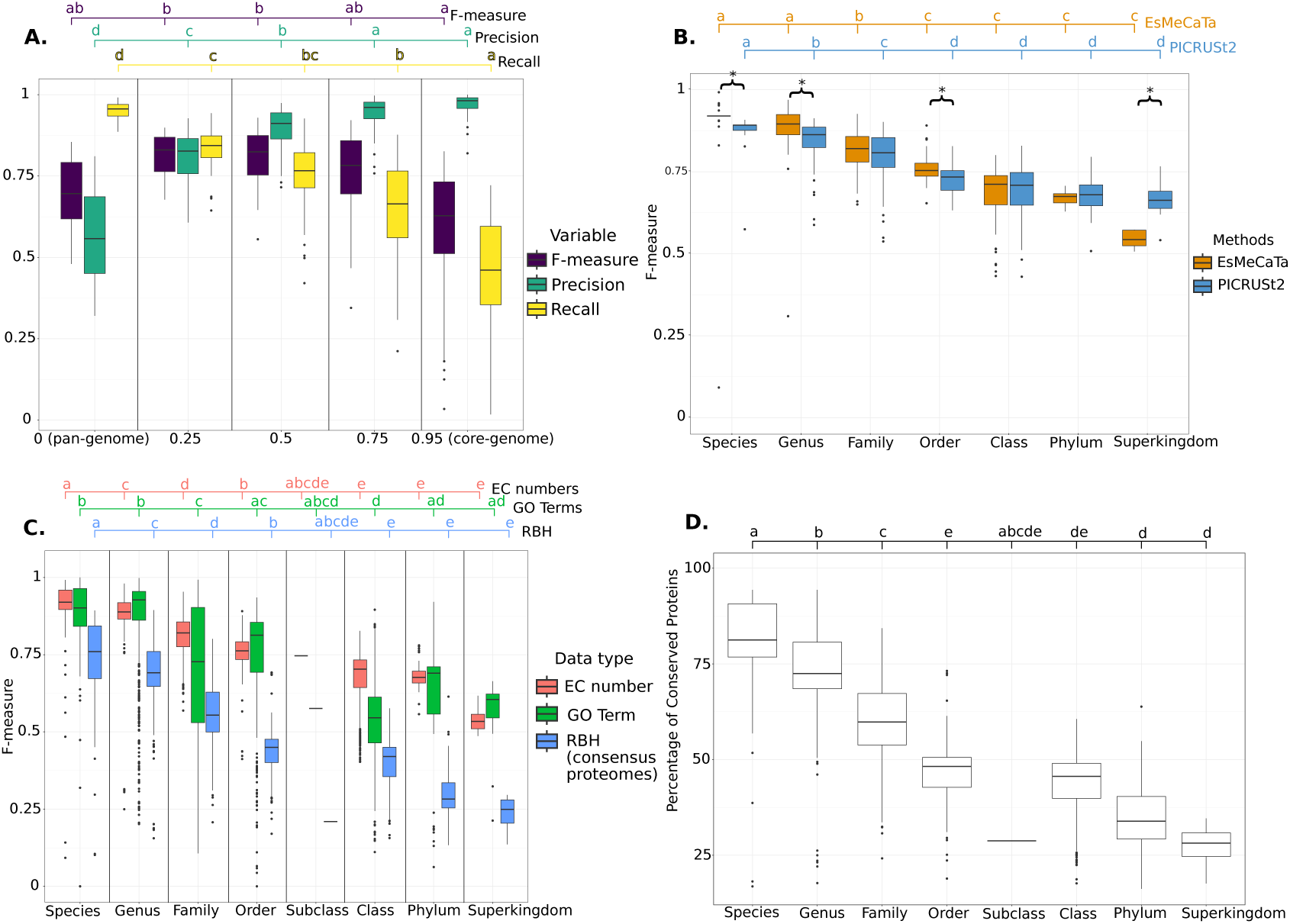
Validation of EsMeCaTa on several datasets. A. Impact of cluster filtering threshold on performance metrics on EC number predictions on the *algal microbial dataset*. F-measures, precision and recall obtained by comparing EsMeCaTa predictions to genome annotations on the EC number predictions. Comparison with 10 paired data of 16S rRNA and complete genomes from [38] and 35 MAGs from [39]. B. Comparison of EsMeCaTa and PICRUSt2 predicted ECs with annotations from MAGs of *MGnify dataset*. F-measures measured on predictions of both methods according to the taxonomic ranks over the whole 565 MAGs. Taxonomic ranks of the MAGs correspond to the rank selected by EsMeCaTa. Bracket with star indicates significant difference between F-measures when comparing the two methods inside a taxonomic rank, according to a Mann–Whitney U test (see Methods). C. Validation of EsMeCaTa predictions against 3,664 MAGS from MGnify database according to the taxonomic rank selected by EsMeCaTa of *MGnify dataset*. F-measures computed from EC number, GO Terms and Reciprocal Best Hits (RBH) between EsMeCaTa proteomes and their associated MAGs (used as the reference). D. Validation of EsMeCaTa predicted proteomes against 3,664 MAGS from MGnify database of *MGnify dataset*. Percentage of Conserved Proteins (POCP) between EsMeCaTa predicted proteomes and their associated MAGs according to the taxonomic rank used by EsMeCaTa to make predictions. Compact Letter Display (a, b, c, d and/or e) indicates significant difference of Dunn’s post-hoc tests when comparing variable distribution (such as F-measures) according to either the threshold used (Panel A) or the taxonomic ranks (panels B, C and D). For more information on compact letter display, see Methods.

### EsMeCaTa performed similarly to PICRUSt2 for EC prediction

MAGs from the MGnify database [5, 6] were used as a reference to estimate the predictive performances of the method. For each MAG of the *MGnify dataset*, four elements were extracted and used in this analysis: (1) their taxonomic affiliations, (2) their proteomes, (3) their annotations (resulting from eggNOG-mapper runs applied to their protein contents) and (4) their predicted rRNAs. The MAG taxonomic affiliations were considered as input to EsMeCaTa, which predicted the corresponding consensus proteome and annotations. These results were compared to the MAG’s own protein sequences and annotations.

Four MAG datasets were considered from diverse environments: honeybee-gut-v1-0 (627 MAGs), human-oral-v1-0 (1,225 MAGs), marine-v1-0 (1,504 MAGs) and pig-gut-v1-0 (3,972 MAGs, see Methods). Among these MAGs, only the ones with a completeness greater than or equal to 90% were kept, reducing these datasets to a total of 3,664 MAGs. Among the 3,664 MAGs, 1,094 MAGs contained 16S rRNA sequences (Table 2), of which 565 could be analysed using PICRUSt2 [8, 9] (198 from honeybee-gut, 55 from human-oral, 187 from marine and 125 from pig-gut datasets). The decrease in MAGs used for this analysis from 3,664 to 1,094 could be explained by the fact that 16S rRNA sequences are often missing from MAGs [40]. The decrease from 1,094 to 565 was caused by poor alignment of the sequences to PICRUSt2 reference sequences (no match with identity superior or equal to 80%).

**Table 2.** Number of MAGs considered for validating EsMeCaTa predictions, of 16S rRNA available in those MAGs and of 16S rRNA that could be analysed using PICRUSt2. Columns correspond to taxonomic ranks selected by EsMeCaTa. The row denoted as "EsMeCaTa" indicates the number of MAGs in the four datasets having a completeness greater or equal to 90% and considered by EsMeCaTa. The row denoted as "16S rRNA" corresponds to the number of MAGs used by EsMeCaTa and having a predicted 16S rRNA. The row "PICRUSt2" corresponds to the number of MAGs with 16S rRNA sequence from which PICRUSt2 was able to make predictions.

| Rank | species | genus | family | order | sub-class | class | phylum | super-kingdom | Total |
| --- | --- | --- | --- | --- | --- | --- | --- | --- | --- |
| EsMeCaTa | 247 | 1,279 | 1,209 | 291 | 1 | 541 | 64 | 32 | 3,664 |
| 16S rRNA | 80 | 422 | 281 | 100 | 1 | 155 | 37 | 18 | 1,094 |
| PICRUSt2 | 39 | 226 | 133 | 50 | 0 | 90 | 19 | 8 | 565 |

EC numbers predicted by PICRUSt2 from these 565 16S rRNA sequences were extracted and compared to the corresponding MAG annotations to compute F-measures. These F-measures were then compared with the F-measures from EsMeCaTa predictions. GO Terms predicted by EsMeCaTa were not used in the comparison as PICRUSt2 did not predict them. The computed F- measures indicate that both PICRUSt2 and EsMeCaTa have comparable performances to predict EC numbers for an organism (Fig 3 B).

The two method performances were further examined according to the taxonomic ranks considered by EsMeCaTa for prediction. Both EsMeCaTa and PICRUSt2 achieved better performances for the lowest taxonomic ranks (such as species and genus, Fig 3 B, Kruskal-Wallis chi-squared = 713.19, df = 6, p-value < 2.2𝑒−16). Low taxonomic ranks considered by EsMeCaTa for predictions imply that closely related proteomes are available in UniProt, which are expected to encompass more similar protein contents and sequence homology, thus helping in accurate comparative genomics predictions. In contrast, higher taxonomic ranks involve larger taxonomic diversity, broader range of gene contents and higher homologous sequence divergence, impeding prediction accuracy.

The decrease of F-measures could be explained by a similar availability and diversity of genome or proteome in UniProt (used by EsMeCaTa) and in the PICRUSt2 database. This suggests that PI- CRUSt2 also has decreasing prediction performances for the organisms from less described taxo-nomic groups. Note however that the taxonomic ranks indicated on the abscissa of Fig 3 B correspond to the ranks used by EsMeCaTa to make the predictions. A similar information is given by PICRUSt2 with the "Nearest Sequenced Taxon Index".

### Accurate prediction of functions until order when compared to MAGs from MGnify

The prediction made by EsMeCaTa on the 3,664 MAGs were assessed regarding three features:(1) EC numbers, (2) Gene Ontology terms and (3) protein contents. For the latter comparison, the consensus sequences of the homologous protein clusters predicted by EsMeCaTa were compared to the corresponding protein sequences of the MAGs (Fig 3 C). Confusion matrices were created and F-measures were computed from them (see Methods).

The results of EC numbers comparison in Fig 3 C presented similar patterns as the comparisons in the previous section with the 565 MAGs. GO Terms showed similar F-measures as EC numbers but with a greater variability. This may be due to the fact that GO Terms include more diverse annotations (metabolism, regulation, localisation) than EC numbers and thus GO Terms are more numerous (around 45,000 terms compared to 9,000 EC numbers), possibly encompassing less conserved annotations.

For all taxonomic ranks, the two functional annotations (GO terms and EC numbers) had better F-measures than the consensus protein sequences. These annotations were predicted from a subset of the proteins. A possible explanation of the difference could be that the functional annotations are inferred from the most conserved protein sequences that are more easily predicted by EsMeCaTa.

A complementary analysis was performed to identify the impact of the dataset on EC numbers, GO Terms and RBH predictions (see Supplemental Figure Sup Fig S2). The honeybee and human oral datasets exhibited better predictions than the marine and pig gut datasets, highlighting the heterogeneity of knowledge available for different environments.

### EsMeCaTa consensus proteomes obtained relevant POCP score at the genus level when compared to MAGs from MGnify

To refine the results from the previous section, another analysis on the consensus proteomes of EsMeCaTa was performed using the Percentage of Conserved Proteins (POCP) metrics [41]. POCP is a metrics used to compute the similarity between two proteomes. It was defined to create boundaries between prokaryotic genera based on protein sequence similarity. A POCP greater than 50% was defined to assign a proteome to a specific genus. In this article, the POCP metrics was considered to estimate the similarity of the consensus proteome of EsMeCaTa to the corresponding MAG. A majority of proteomes estimated from genus rank had POCP ranging from 50% to 90% (Fig 3 D). These values are close to the ones proposed by [41], supporting the fact that the consensus proteomes estimated by EsMeCaTa, in term of sequence conservation, could be considered as belonging to the same genus as their associated MAGs. Thus, it could be used as a representative of the MAG taxonomic group. As expected, this metrics decreases as the taxonomic rank increases (Fig 3 D). Overall, in terms of conserved proteins, these results suggest that EsMeCaTa provides accurate estimates of proteomes up to the genus and family ranks.

### Exploring predicted functions in a methanogenic reactor microbial community

The microbial community from an experimental methanogenic reactor was studied using a metabarcoding sequencing approach. Methanogenic reactors are anaerobic biological processes where microbial communities (composed of bacteria and methanogenic archaea) degrade complex organic matter mainly into methane, CO_2_ and water. During a 195 days long experiment, 27 samples of digestate were retrieved from the reactor and their microbial communities were characterised by DNA extraction and 16S rRNA high throughput DNA sequencing (see Methods and Supplementary materials). Reads were analysed using FROGS [42], providing taxonomic affiliations for 445 operational taxonomic units (OTU, see Methods). For each time point, OTU relative abundances (Sup Fig S3), biogas production and additional physico-chemical parameters (Sup Fig S4) were measured.

The diversity of the OTUs was in line with previous studies, exhibiting a community dominated by the phyla *Bacillota* (formerly *Firmicutes*, see [43]) and *Bacteroidota* (formerly *Bacteroidetes*, see [43]), representing 53% and 16% respectively of the OTUs [44, 45] (see Supplementary Materials and Sup Fig 3). Few affiliations were as precise as species (58), most corresponded to the ranks genus and family (306), and 81 were assigned to taxonomic ranks higher than order (lines in Sup Fig S5).

The 445 taxonomic affiliations were used as input to EsMeCaTa. The method selected the taxonomic ranks suitable for prediction according to the proteomes availability in UniProt, providing insights into the knowledge available for the studied community (Fig 4 A, generated by *esme- cata_report* command). For the 364 OTUs identified at the species, genus or family rank by FROGS, 79% (289) were selected by EsMeCaTa at a taxonomic rank of species, genus or family (Sup Fig S5). For the other OTUs, EsMeCaTa considered higher taxonomic ranks, such as class (72 OTUs) or phylum (44). This indicated the presence within the methanogenic community of organisms that are weakly characterised at the genomic level [44]. This concerned, for example the presence of OTUs from understudied bacterial lineage, such as phyla *Cloacimonadota* [46] or *Hydrogenedentes* [47, 48].

**Figure 4.**
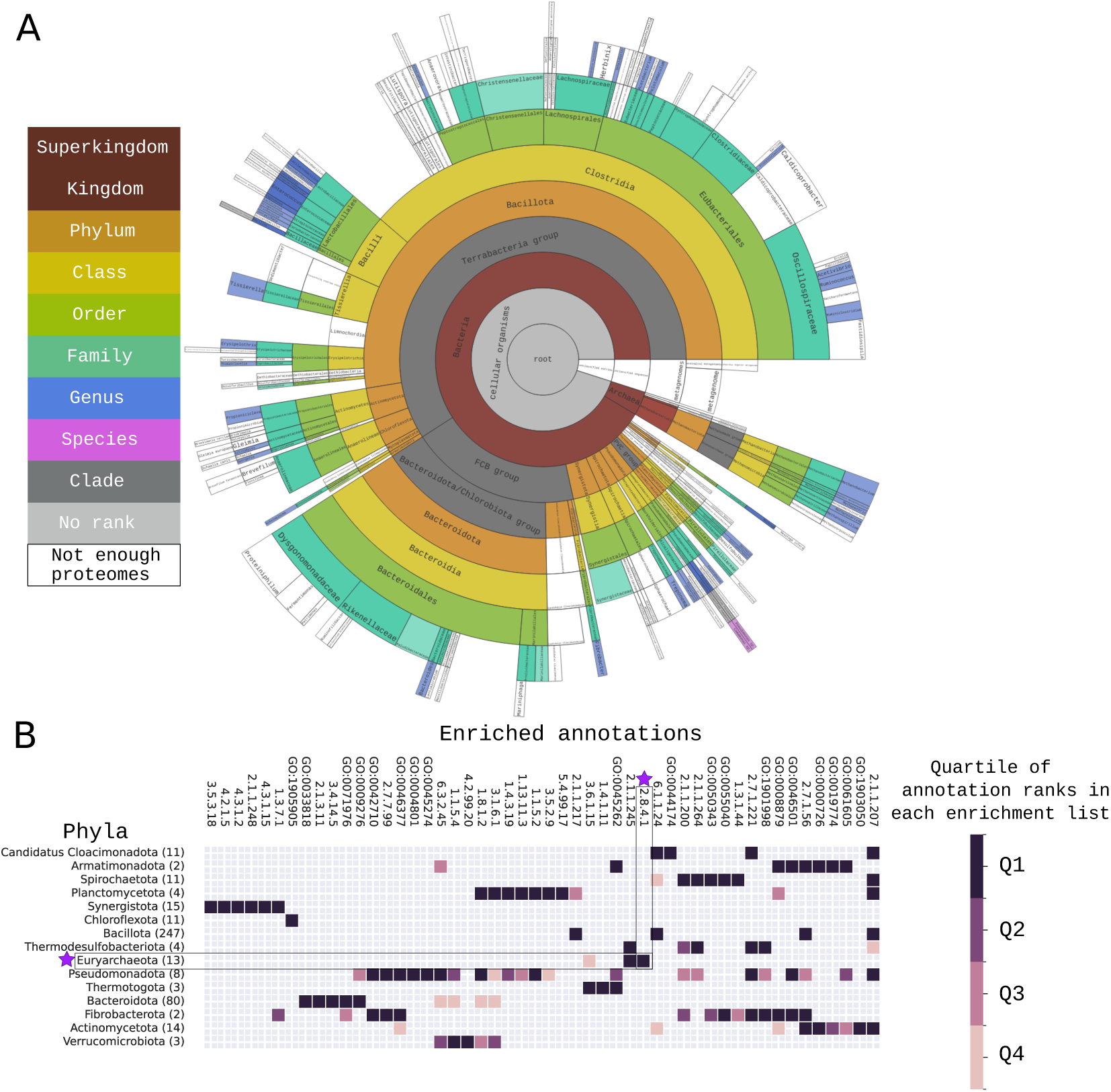
A. Methanogenic reactor taxonomic diversity. Taxonomic diversity of EsMeCaTa selected taxa for the 445 OTUs estimated from 16S rRNA amplicon sequencing of an experimental methanogenic reactor. Colored taxonomic boxes correspond to taxon selected by EsMeCaTa. Empty boxes indicate the absence of enough proteomes and the need of EsMeCaTa to use a higher taxonomic rank. **B.Annotations (GO terms and EC numbers) enriched in the methanogenic reactor major phyla, compared to the whole community annotations.** The figure is produced using Orsum [37]: the enriched functions (GO Terms and EC numbers) are shown on the left figure side, coloured squares in the middle matrix indicate the taxa owning the enriched functions, and the colour scale on the right figure side indicates the enriched function rank, estimated with GSEApy [36] and binned into quartiles. Number of OTUs contained in the phyla are indicated in parenthesis after the phylum name. Purple star indicates the focus on the Euryarchaeota phylum and the EC number 2.8.4.1 associated with final step of methanogenesis.

Thousands of metabolic functions were predicted by EsMeCaTa for this methanogenic community (Sup Figure S6 A and B). An enrichment analysis of metabolic functions across phyla was first performed using GSEApy [36] and Orsum [37] in order to automatically highlight the major differences between the phyla present in the community (see Methods). Among the identified functions, two of the three functions enriched in the phylum Euryarchaeota were associated with methanogenesis. These corresponded to the EC number 2.8.4.1 catalysing the last reaction of methanogenesis and the EC number 2.1.1.245, a key enzyme in methanogenesis from acetate (Fig 4 B and Sup Fig S7, generated by *esmecata_gseapy* command). This is congruent with the diversity of the Archaea identified in the reactor, which are specifically methanogens (Fig 4 A), and with the fact that methanogenesis in reactor conditions is exclusively performed by archaeal species [49].

But not all functions could be searched with simply functional annotations. For example, cellulose degradation performed by cellulosome is expected in methanogenic reactor. This intricate complex is, for example, described by the MetaCyc database as a multi-step reaction (reaction RXN-14887) without EC number assignment. A method has been proposed to identify potential organisms containing them by using Genomics and Bioinformatics tools [50]. It is based on homology search and it has been applied to EsMeCaTa consensus proteomes in order to identify taxon containing cellulosome in the samples. Reference dockerin and cohesin sequences from the literature were aligned using Diamond [51] against the consensus proteomes (see Method). Both proteins matched consensus sequences predicted for genera *Acetivibrio* and *Ruminiclostridium* (Sup Fig S8). This suggests a cellulosome activity in those taxa, in agreement with previous works [52, 53].

### Characterising Archeae trophic groups operating methanogenic pathways in the biore- actor

Several methanogenic pathways were profiled using EsMeCaTa predictions. An overview of these pathways was generated by combining pathways from metabolic databases and literature (Fig 5, see Methods). The pathway reactions were then searched in two ways : either by directly identifying EC numbers in the function table, or by aligning reference sequences from SwissProt or KEGG Orthologs to the consensus proteomes (see Methods).

**Figure 5.**
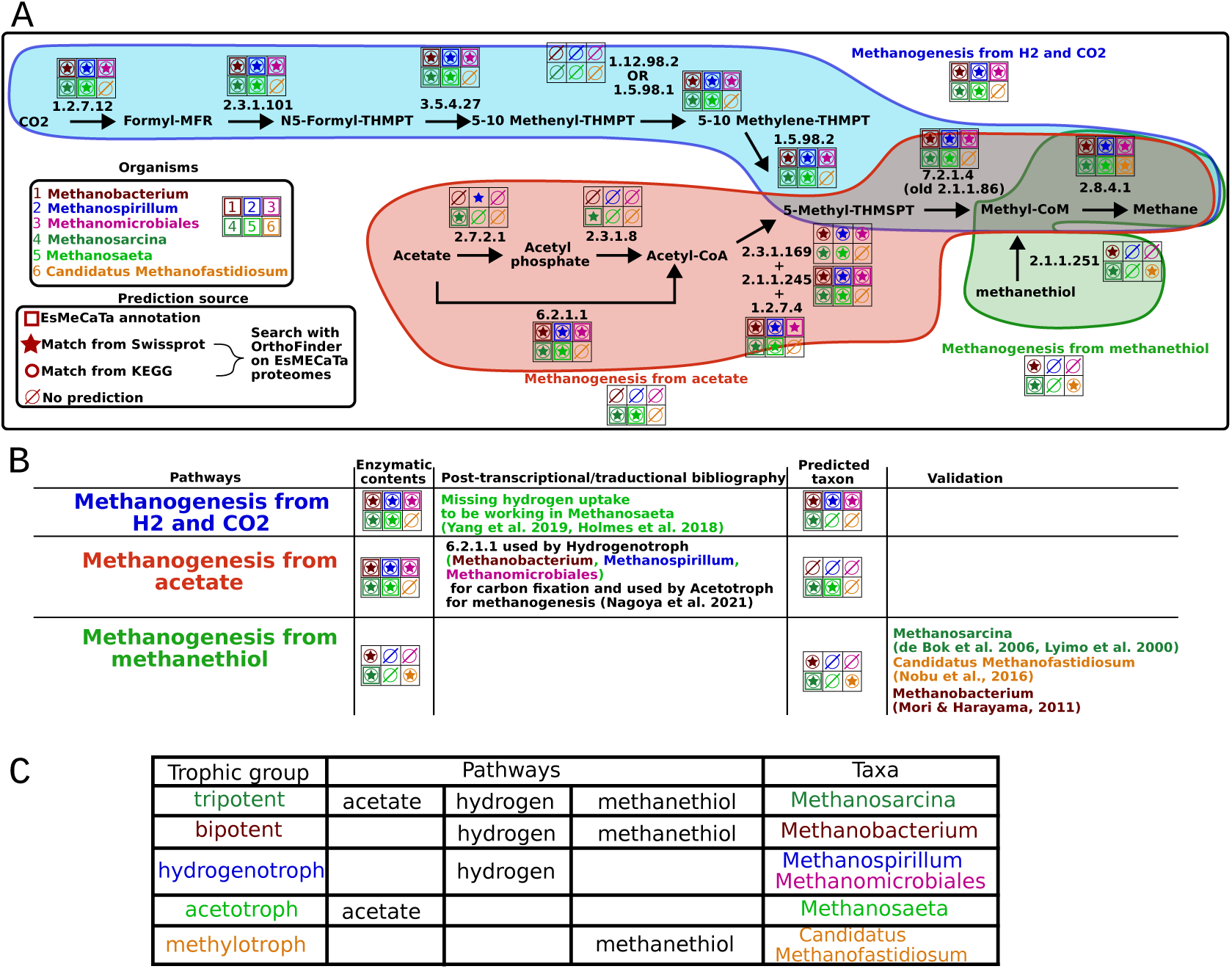
Enzymatic characterisation of three metabolic pathways for methanogenesis. A. Schematic representation of pathways producing methane from CO_2_, acetate or methanehtiol. Square, star and circle indicated if the enzymes was found in the corresponding organism. Enzymes identified from EsMeCaTa annotation were shown with square. For enzymes associated with SwissProt or KEGG Orthologs, alignment made with Diamond [51]. Results were filtered according to RBH procedures. If an enzyme matched a sequence of SwissProt associated with an EC, it was shown with a star and if it matched with a KEGG Orthologs, it was shown as a circle. **B. Table showing literature reference for each pathways. C. Table showing the different identified groups.** Each OTUs were put in different trophic groups.

First, reactions from the hydrogenotrophic methanogenic pathway were found in all Archaea of the community (by the three annotation procedures), except in *Candidatus Methanofastidiosum* (Fig 5). This is expected as most of the methanogenic archaea observed in reactor conditions are known to perform hydrogenotrophic methanogenesis [44]. Note however that the presence of the enzymes does not necessarily indicate that the pathway is active in the community: *Methanosaeta* was predicted to possess all the required enzymes, but it can not use hydrogen as an electron donor and relies on a direct interspecies electron transfer [54, 55].

Second, concerning the acetotrophic methanogenic pathway, EC numbers 2.7.2.1 and 2.3.1.8 were found only in *Methanosarcina* (Fig 5), consistently with previous results showing its ability to perform both hydrogenotrophic and acetotrophic methanogenesis [56]. An alternative reaction involves EC number 6.2.1.1 for acetate degradation, which is used for methanogenesis specifically in *Methanosaeta* [57, 58]. This reaction was found in all Archaea of the community, except *Candidatus Methanofastidiosum* (Fig 5). These reactions are reversible and involved in carbon assimilation through acetate synthesis [59], such as performed by the Wood–Ljungdahl pathway (WLP). Congruently these reactions were predicted in a wide range of bacterial clades, with some known to perform WLP (Sup Fig S9 and S10). The next step of the acetotrophic methanogenic pathway, EC number 2.3.1.169, was not predicted by EsMeCaTa annotation step but it was found with alignment to sequences from SwissProt and KEGG Orthologs.

Third, for the methanogenic pathway from methanethiol (Fig 5), EC number 2.1.1.251 was found only in the predicted annotations for *Methanosarcina*, which is consistently known to achieve this pathway [60, 61]. Moreover, using sequences homology searches, the reaction was also found in *Candidatus Methanofastidiosum* (which was expected according to literature [62]) and in *Methanobacterium* (which contains species that perform this function [63]).

Altogether, three methanogenic pathways were profiled in six taxa using EsMeCaTa annotations, consensus proteomes with subsequent annotation procedures and literature validations (Fig 5). These results illustrated also how pathway profiling can be achieved using EsMeCaTa predictions. In particular the absence of predictions for EC number 2.3.1.169 by eggNOG-mapper showed the limit of relying on specific annotation, and the usefulness of consensus proteomes for additional homology search.

The EsMeCaTa annotations and consensus proteomes predictions enabled the association of the three methanogenic pathways with OTUs exhibiting the enzymatic potential characteristic of these pathways (Fig. 5 B). Our method predicted that four taxa can activate hydrogenotrophic methanogenesis (associated with 11 OTUs), two taxa can activate acetotrophic methanogenesis (associated with three OTUs), and three taxa can activate methylotrophic methanogenesis (associated with eight OTUs).

Since each pathway is linked to a specific substrate and to several organisms, OTUs were grouped into different trophic categories based on the number of substrates they are predicted to degrade. Five trophic groups were identified in the bioreactor (Fig. 5 C), including complex trophic groups capable of activating multiple pathways, such as *Methanosarcina*. More specifically, the tripotent group comprises two *Methanosarcina* OTUs; the bipotent group (having both hydrogenotrophic and methylotrophic pathways) includes five *Methanobacterium* OTUs; the hydrogenotrophic group consists of four *Methanospirillum* or *Methanomicrobiales* OTUs; the methylotrophic group contains one *Candidatus Methanofastidiosum* OTU; and the acetotrophic group contains one *Methanosaeta* OTU.

The five trophic groups include all the methanogenic taxa identified in the community by EsMeCaTa. Thus we hypothesised that the biogas production could be explained by these organism abundances. In the next sections we investigate the dynamics of these groups in light of the biogas production over time.

### Linking detected functions to microbial groups abundances

To assess the impact on biogas production of the five trophic groups associated with methanogenesis, as predicted from the EsMeCaTa output (Fig. 6 A), we compared the relative abundances of these groups to biogas production. For each trophic group, the relative abundances of their respective OTUs were summed and then normalized using z-score normalization (see Methods). The normalized abundances were subsequently analyzed in relation to bioreactor perturbations caused by different intakes over time (pig slurry phase, fruit phase, fat phase, and food waste phase).

**Figure 6.**
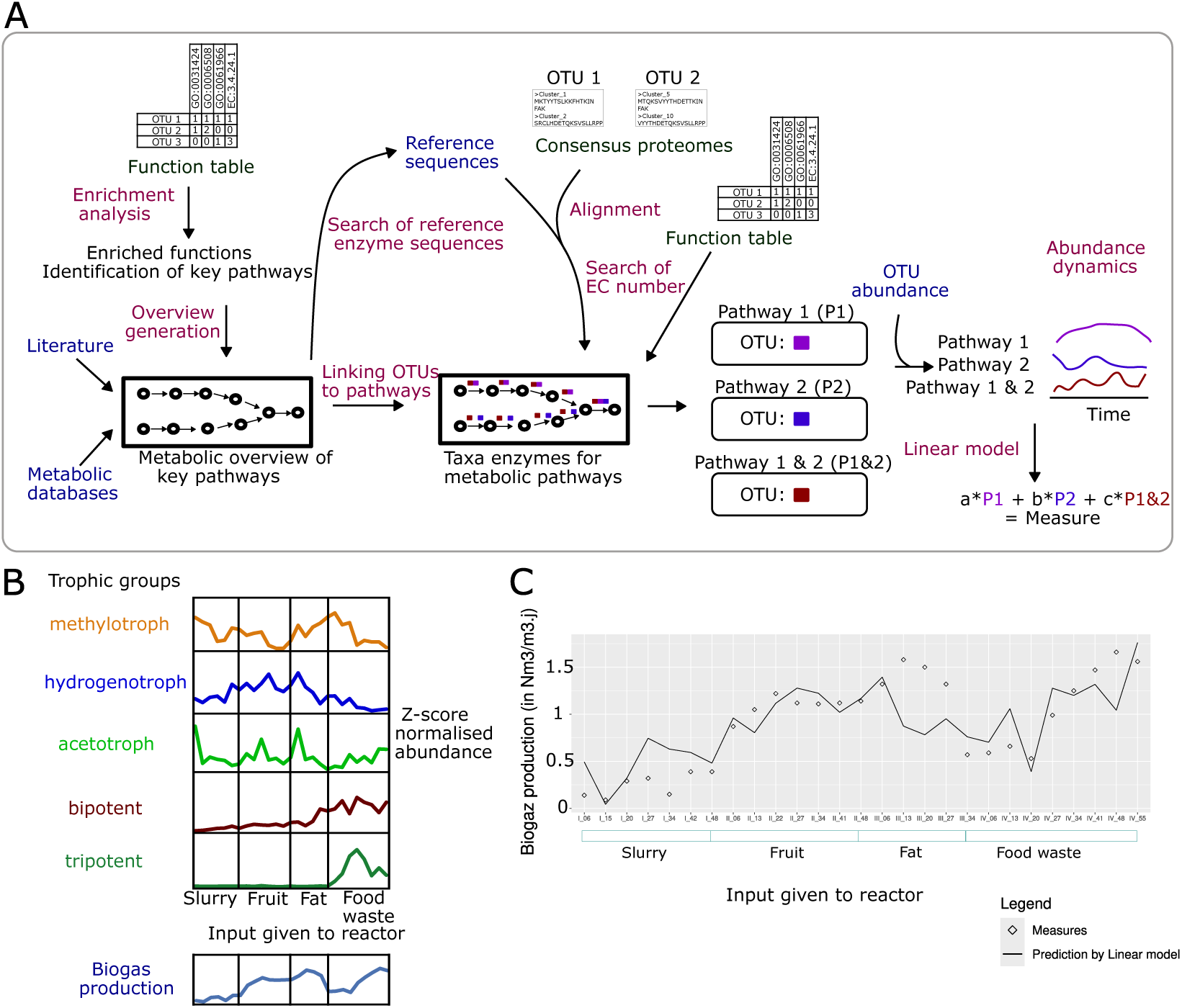
A. Workflow analysis of methanogenic pathways. Steps to the identification of the different methanogenic pathways according to the abundance of the associated OTUs. First, using EsMeCaTa function table, enriched annotations in phyla were identified. Among them, the final EC number of methanogenesis was found to select OTU associated with it. Then using literature on the associated taxa and metabolic databases, an overview of methanogenic pathways was generated. This overview was completed by finding EC number present in the taxa thanks to EsMeCaTa function table and homology search with the consensus proteomes. **B. Dynamics of abundance of groups.** For each trophic group, abundance from their OTUs was summed and normalised with a z-score normalisation to look at their dynamics over time points and according to perturbation (slurry, fruit, fat and food waste intake. **C. Linear model prediction.** Comparison of biogas production with predicted biogas from a linear model made from the abundance of the different trophic groups.

As shown in Fig. 6 B, the time series of trophic group abundances exhibited distinct and characteristic behaviors, and none of them clearly correlates with the biogas production measurements. The methylotrophic group was abundant during the slurry and fat phases. For the slurry fate, this is consistent with the presence of methanethiol in slurry [64]. The hydrogenotrophic group was particularly abundant during the fruit and fat phases, whereas the acetotrophic group displayed a peak in abundance at the beginning of the slurry phase and increased during the fruit and fat phases. Both the tripotent and bipotent groups showed increased abundances when food waste was added.

To test the cumulative effects of the different trophic groups on biogas production, we applied a linear model (see Methods), whose predictions are shown in Fig. 6 C. The model predicted that the combined abundances of Archaea explained biogas production significantly better than the intercept alone (F-statistic: 6.018 on 5 and 21 degrees of freedom, p-value = 0.001321). Two groups were identified as key contributors to this result: the bipotent group containing Methanobacterium (p-value = 0.000588) and the methylotrophic group containing *Candidatus Methanofastidiosum* (p-value = 0.001781).

These findings suggest that the trophic groups identified through post-processing of EsMeCaTa outputs are sufficient to statistically and significantly predict biogas production under a cumulative hypothesis. This analysis underscores the importance of performing multiple analyses using EsMeCaTa results and combining them with post-analyses (as shown in Fig. 6 A) to better understand complex behaviors within a bioreactor subjected to multiple perturbations.

## Discussion

In this article, we described a method to predict protein sequences and functions for taxa from their taxonomic affiliations. The method was applied to several datasets in order to validate the predictions and to illustrate how they might be considered for further investigations. By giving additional information on the prediction (selected taxonomic rank, available proteomes, consen-sus protein sequences associated with the predictions), EsMeCaTa gives an explainable way to assess the predictions and filter them. These results can be both automatically analysed, for example using enrichment analysis, or manually investigated as shown in the section focusing on the methanogenic reactor.

### Handling the heterogeneity of the sequencing data characterizing the environmental taxonomic diversity

Several alternative phylogenetic marker genes are also considered to study environmental communities, such as 18S rRNA for eukaryotes [65], ITS for fungi [66], or *gyrB* [67] and *rpoB* [18] genes for bacteria. Obtaining taxonomic affiliation from these gene amplicon sequences is made possible thanks to plethora of methods [68]. Furthermore, other approaches can be considered to profile the environmental taxonomic diversity, such as *shallow whole genome sequencing* [19, 20], metatranscriptomics [21] or long read sequencing [69].

EsMeCaTa has been designed to handle these numerous heterogeneous technologies for metabarcoding and metagenomics, by taking as input the common predictions issued from these data : the sequenced reads’ taxonomic affiliations. We demonstrated this flexibility by applying EsMeCaTa to several datasets: (1) an example containing 13 manually selected taxa ranging from genus to class, (2) taxonomic affiliations of MAGs and complete genomes from metagenomics, and (3) taxonomic assignment from metabarcoding of 16S rRNA genes. Thus EsMeCaTa appears suitable to compare predictions at a functional level issued from different sequencing technologies.

As a perspective, EsMeCaTa could also be used to link metabarcoding and metagenomics in the same experimental study, especially in time-series community measurements that combine a large number of metabarcoding samples with a few metagenomics samples. In such an experimental setting, EsMeCaTa could be configured to use protein predictions from the assembled metagenomes as a basis for function prediction instead of UniProt, at least for the taxa represented in the metagenome. Thanks to this pairing, the predicted metagenomic profiles could thus incorporate the particularities of the genomes of the local species, while benefiting from the higher spatial and temporal resolution provided by metabarcoding approaches.

Similarly, EsMeCaTa could be applied to the analysis of culturomics data banks. This high-throughput culture approach combines the taxonomic characterization of the bank through amplicon sequencing and the complete genome sequencing of a few selected organisms [70]. In this context, EsMeCaTa could expand functions associated with complete genomes with functional predictions related to all the bank’s organisms.

### Highlighting bias due to the heterogeneity of knowledge associated with taxa

EsMeCaTa predictions were compared to protein sequences and annotations of MAGs from the MGnify datasets and with annotations from paired data consisting in 16S rRNA sequences and complete genomes or MAGs for an algal microbiota dataset. With these comparisons, we illustrated the impact of available knowledge (here proteomes from UniProt) according to the taxon used as input. The quality of these predictions were shown to be dependent on the taxonomic ranks that were selected by EsMeCaTa, less available knowledge requires the use of more distant organisms and to select larger taxon, then impeding the predictions.

In a comparison with PICRUSt2 on the MGnify datasets, we showed that both methods have similar performance for EC predictions and that both methods were impacted by the issues of knowledge availability (despite not using the same database). This highlighted the impact of uncertainty and the explainability given by such methods on functional estimation.

A complementary explanation to the loss of quality prediction of functional annotation methods such as EsMeCaTa and PICRUST2 is the ecology and adaptation of the organisms present in the taxon selected by the tools. Indeed, ecological diversity of organisms in a taxon impacts the pan- genome of taxon by the number of shared and unique genes. This is the case for open pan-genome when newly sequenced genomes continuously reveal new genes [32]. This is for example the case with the open pan-genome of the species *Escherichia coli* [71]. In contrast "closed" pan-genomes denote taxa encountering few horizontal gene transfers. This highlights the difficulty for any tool to estimate organisms potential in taxa with open pan-genomes, due to the potential unique genes present in these organisms resulting, for example, from the numerous horizontal gene transfers providing new and rare functions.

### Explaining and refining predictions with consensus proteomes and intermediate information

In order to give more insights on the step leading to the prediction of function, EsMeCaTa provides intermediary information such as the taxon used by EsMeCaTa, proteomes used, consensus proteomes estimated… As demonstrated with the methanogenic reactor dataset, this allows for a better understanding of the predictions made by the method but also to make advanced search. Consensus proteomes, in particular, are an insightful result of EsMeCaTa that was leveraged to explore the protein potential of a taxon, for example, by searching specific databases or by characterizing protein complexes.

More generally, by relying on UniProt IDs, EsMeCaTa provides a link between microbial community datasets and either cross-references from many other databases or results, such as the predicted structure for millions of proteins [27, 72]. Another information of interest is the environmental conditions associated with the studied organisms. Currently EsMeCaTa retrieves all the proteomes of a taxon. Among the selected proteomes, some could be linked to organisms that could not be living in the environmental conditions studied. Thus this could require a new filtering step according to the known living conditions of the associated organisms, for example pH, temperature aerobic or anaerobic. This information could be retrieved from databases, such as the BacDive database [73]. As perspective, we plan to use this information both to increase the prediction accuracy of EsMeCaTa and to filter the proteomes used to estimate the consensus proteomes.

### Taking knowledge advances into account

A key feature of the EsMeCaTa is to be up to date with the latest knowledge available from UniProt, giving the advantage of computing consensus proteomes associated with newly identified taxa and following updates from the taxonomy database. But this comes with a cost on performance (Sup Table 1), reproducibility related to updates of UniProt or NCBI Taxonomy databases, impact on UniProt servers and ecological impact (necessity of downloading and computing for each run launched by users). As another perspective, we plan to improve EsMeCaTa by creating precomputed database of proteomes predictions according to new release of UniProt associated with specific version of the NCBI Taxonomy database. This will require to parse most of the taxa present in the UniProt database and apply EsMeCaTa on these taxa to create the precomputed database. Relying on this precomputed database would speed up the predictions and avoid the reprocessing of the different taxa.

### Shifting from population abundances to individual taxa

Many functional characterization methods include a final step of functional profiling based on abundance data. However, most are tailored to specific gene markers and may be considered challenging [74, 75]. For example, when functional profiles are created using 16S rRNA gene sequencing, estimating the functional abundances that would have been measured with metagenomic data requires several steps. These include weighting predicted functions by OTU abundances and, in the case of 16S rRNA amplicons, normalizing them by gene copy numbers [8, 9].

In this work, however, given that EsMeCaTa accepts a wide range of inputs from different sequencing technologies, the main pipeline does not include functional profiling based on organism abundances. Instead, we adopted a post-processing strategy to enrich functional tables with taxon abundances and generate comprehensive functional profiles tailored to specific contexts.

The first example of such post-processing is illustrated in the case study on methanogenesis. Here, functional annotations predicted by EsMeCaTa were used in a post-analysis to design func-tional profiles for trophic groups associated with methanogenic pathways. Specifically, we calculated the sum of OTU abundances involved in each trophic group. Finally, a z-score normalization was applied, enabling comparison of the trophic group abundance dynamics with methane production over time. This approach facilitated an assessment of how trophic group dynamics correlated with methane production trends.

A second example of post-processing for functional profiling is described in [76]. In this study, EsMeCaTa was applied to metagenomic datasets to associate taxa with functions, followed by a machine learning approach to identify discriminative functions. Function abundance values were calculated using a two-step procedure: first, for each taxon in a sample, the number of protein clusters associated with a given function was summed and multiplied by the taxon’s abundance. Then, across all taxa in the sample, these values were summed to obtain the total abundance for each function. These computed function abundances were subsequently used to train random forest classifiers to distinguish between patient and healthy individuals. Interestingly, classification based on predicted function abundances achieved comparable performance to classification based on organism abundances, while revealing hidden cumulative effects in microbiomes. As a future direction, we anticipate adapting the classification strategy developed in [76] to multi-level data, which could provide additional insights into methanogenesis dynamics.

## Conclusion

EsMeCaTa is a new software to estimate consensus proteomes and metabolic functions from taxonomic affiliations. To handle results from different sequencing approaches (metagenomics or metabarcoding), EsMeCaTa relies on the taxonomic affiliations inferred from the sequencing data. This software provides several intermediary results to help understanding its predictions and allowing users to make additional analyses and annotations thanks to predicted consensus proteomes. Benchmark between EsMeCaTa predictions and MAGs to exhibit its predictions. Furthermore, EsMeCaTa and PICRUSt2 were compared to show their similar performances. The possibility of EsMeCaTa to study metabarcoding data was shown using a novel dataset from a methanogenic reactor. This software gives a flexible method to study microbial communities from environmental data.

## Methods

### Datasets

Four datasets were considered in the article: (1) an arbitrary taxa list, (2) a dataset of MAGs and complete genomes from symbiotic bacteria of *Ectocarpus sp.* brown algae (3) a dataset of Metagenome-Assembled Genomes (MAGs) from the MGnify database and (4) a microbial community sequenced from an experimental methanogenic reactor.

### Benchmarking datasets

The *arbitrary taxa list dataset* contains thirteen taxa manually selected to illustrate the EsMeCaTa workflow and its main outputs. Taxa were separated into two groups, a first one containing bacteria close to the *Escherichia* genus and a second one containing eukaryota related to the *Plasmodium* genus. Both these genera contain model species for which multiple proteomes were available, allowing functional prediction at the genus level. Few or no proteomes are associated with the other genera considered, precluding predictions at the genus level. This dataset is available as a list of taxonomic affiliations ().

The *Ectocarpus sp. microbiota dataset* was used to investigate the impact of the cluster filtering threshold, or representativeness 𝑅, on the predictions. It contains 10 paired data associating each a 16S rRNA gene sequence and a complete genome from [38, 77] and 35 MAGs from [39], having a completeness greater than 90%. The taxonomic affiliations for the complete genomes were obtained from the 16S rRNA sequencing of the associated organisms available in [77]. The taxonomic affiliations for the MAGs were extracted from the supplemental Table S4 in [39]. The 45 taxa affiliations are provided in the The *MGnify dataset* was obtained from the MGnify database [5, 6] and used to estimate the accuracy of the predicted functions and consensus protein sequences. The following genome catalogues of MGnify were used: honeybee-gut-v1-0 (627 MAGs, taken from https://ftp.ebi.ac.uk/pub/ databases/metagenomics/mgnify_genomes/honeybee-gut/v1.0/), human-oral-v1-0 (1,225 MAGs, taken from https://ftp.ebi.ac.uk/pub/databases/metagenomics/mgnify_genomes/human-oral/v1.0/), marine- v1-0 (1,504 MAGs, taken from https://ftp.ebi.ac.uk/pub/databases/metagenomics/mgnify_genomes/marine/v1.0/) and pig-gut-v1-0 (3,972 MAGs, taken from https://ftp.ebi.ac.uk/pub/databases/metagenomics/ mgnify_genomes/pig-gut/v1.0/).

The subset of 3,664 MAGs with a completeness greater than or equal to 90% were considered for benchmarking ().

### Methanogenic community case-study

The *methanogenic reactor dataset* corresponds to the diversity sequenced from an experimental methanogenic reactor and illustrates the application of EsMeCaTa to a real metabarcoding dataset. A detailed description of the experimental setup was published in [78] and is summarized in the section "Summary of the methanogenic reactor operation and microbial community characterization". The sequencing procedure is described below.

Digestate was sampled weekly from the methanogenic reactor for 195 days to perform a total DNA extraction of its microbial community using the NucleoSpin® Soil DNA extraction kit (Macherey-Nagel, USA). Metabarcoding targeted the archaeal and bacterial hypervariable V4-V5 regions of the 16S rRNA genes using the so-called universal primers 515F (5’- Ion A adapter-Barcode-GTGYCAGCMGCCGCG 3’) and 928R (5’-Ion trP1 adapter-CCCCGYCAATTCMTTTRAGT-3’) and PCR amplification. The resulting amplicons were purified, quantified and sequenced at the metagenomic platform of the UR1461 PROSE of INRAE (Antony, France) according to manufacturer’s instructions, and as described in [79]. Sequencing was performed on an Ion Torrent Personal Genome Machine using Ion 316 Chip V2 (Life Technologies) and Ion PGM Hi-Q View Sequencing Kit (Life Technologies).

A total of 2,164,633 raw reads were sequenced from the 27 digestate samples and were processed with the FROGS pipeline [42] following the authors’ recommendations on the MIGALE Galaxy instance (INRAE, Jouy-en-Josas, France). The first processing steps included primer trimming and quality control, resulting in 1,145,396 reads of approximately 380 base pair length without N. The next steps consisted in sequences clustering, chimera removal, low abundance OTU filtering at 0.01%, and taxonomic affiliation of the OTUs with 16S SILVA Pintail100 [65]. It resulted in 1,031,447 clean sequences affiliated to 445 taxa, with a mean of 38,202 +/- 17,300 sequences per sample. The 445 taxa dataset is provided in the . The metabolites measured in the methanogenic reactor are provided in .

### Metadata of the run

All these different run of EsMeCaTa were done using 10 CPUs and 60 GB of RAM. The runtimes taken by the method can be seen in Sup Table S1. The run were performed with EsMeCaTa version pre-release 0.5.0. The different metadata on the version of the used dependencies are present in Sup Table S2.

### The EsMeCaTa workflow

EsMeCaTa is a Python package predicting protein sequences and functions from taxonomic affl- iations that can be called with the command *esmecata*. The step of this pipeline are described in Figure 1 and below. It takes as input a tabulated file containing a list of taxonomic affiliations and it outputs consensus proteomes and functions tables indicating the occurrence of functions (EC numbers and GO Terms) in the taxa. It relies on several Python packages (Biopython, bioservices, ete3, pandas, requests), NCBI Taxonomy database, UniProt database, MMseqs2 and eggNOG-mapper.

#### Identification of available proteomes and selection of associated taxon from taxonomic affiliations

The *esmecata proteomes* command uses the ete3 Python package [80] to parse the input taxonomic affiliation file and assign a NCBI taxon ID [26, 81] to each input taxon. A maximum taxonomic rank can be parameterized by the user to limit the analysis to the lower ranked taxa (option *rank limit*). The complete lineage information is used to determine the appropriate NCBI taxon ID when ambiguities occur.

REST API queries to the UniProt [27] proteome database extract the identifiers of all Uniprot proteomes associated with each NCBI taxon ID. Metadata of the proteomes allows the selection of the taxon ID with the lowest rank in the taxonomic affiliations such that it is associated with at least 𝑁 proteomes (defaut value 𝑁 = 5) having a BUSCO score [31] greater than 80% and not tagged as "redundant" and "excluded" in Uniprot.

For each NCBI taxon ID, if the number of selected proteomes is greater than a parameterised threshold (100 by default), a sub-sampling procedure is performed. A taxonomic tree is created with the input taxon as root and the organism IDs associated with each of the proteomes as leaves. This allows sub-sampling 100 random proteomes which conserves the distribution of proteomes in each sub-group of the taxon and thus the taxonomic diversity. Due to the randomness of the selection during sub-sampling, one can expect variations with the taxon impacted by this procedure. All selected proteomes are then downloaded. The *esmecata check* command performs the same steps without downloading the proteomes, thus simply showing the proteome availability in the Uniprot proteome database.

### Estimation of the consensus proteome from protein clustering and cluster filtering

The *esmecata clustering* command computes a consensus proteome by identifying the proteins shared by the proteomes associated with a taxon. MMseqs2 [28] performs protein sequence clustering from the proteomes and generates consensus protein sequences for each cluster using the most frequent amino-acid at each position of the profile. The sequence identity threshold is chosen to match distantly related homologues, with a minimum sequence identity of 30% and a minimum coverage of 80% [82]. The resulting clusters of homologous proteins are then filtered according to the distribution of the proteins among the proteomes. The representativeness ratio 𝑅_𝑝_ between the number of proteomes represented in a protein cluster and the total number of proteomes selected by the *esmecata proteomes* command is calculated for each cluster. Then the algorithm selects all the protein clusters that contain proteins from at least half of the taxon proteomes, that is 𝑅_𝑝_ ≥ 0.5 with a threshold 𝑇_𝑟_ = 0.5. Other cluster filtering thresholds 𝑇_𝑟_ can be defined by the user. For a given taxon, the set of consensus sequences from the selected clusters is denoted as the *consensus proteome*.

### Creation of the function table and the PathoLogic files from consensus protein annotations

The *esmecata annotation* command uses the consensus proteomes to predict the functions associated with each taxon. Each consensus protein sequence is annotated using eggNOG-mapper [29, 30] with default parameters. The *function table*, which constitutes EsMeCaTa functional predictions for the input taxa, is constructed by counting the occurrence of Enzyme Commission numbers and Gene Ontology terms predicted for each input taxonomic affiliation. This information is also used to create PathoLogic format files, the input format used by Pathway Tools [83, 84] for draft metabolic networks reconstruction.

## EsMeCaTa output post-analysis

### Visual summary of EsMeCaTa results

The *esmecata_report* command produces a HTML report summarising the main predictions of the EsMeCaTa run, created with DataPane (https://github.com/datapane/datapane). The report is divided into panels with figures showing the results of each step of the workflow (drawn with Plotly [85]). The first panel shows the taxonomic diversity of the input taxa and the taxa considered for prediction. A sunburst chart indicates the names of the taxonomic affiliations provided as input, and the taxa selected by EsMeCaTa for predictions (see Fig 4A for an example). A Sankey diagram represents the same information, indicating which input affiliations correspond to which taxa selected for prediction (*i.e.* Fig 2 A). The second panel, *proteomes summary*, shows a summary of the proteome downloading step: the number per taxonomic rank of taxa provided as input and of taxa selected for prediction, and the distribution of the proteome number per taxon. The third panel, *clustering summary*, shows information about the protein clustering step. It displays the number of protein clusters obtained according to the clustering threshold 𝑅, allowing visualization of the pan-proteome distributions (*i.e.* Fig 2 B). The fourth panel, *annotation summary*, shows the amount and categories of EC numbers and GO terms predicted for a given dataset. Such figures illustrate the functional capacity and redundancy from the individual taxon level to the community level. The fifth panel displays summary results for all the previous steps.

For the sake of reproducibility, the last panel displays metadata concerning the used parameters and the dependencies’ versions of the EsMeCaTa run. The report consists in a static HTML file containing all the figures. Each figure is also exported in the user-specified output directory, in HTML for visualization outside the report, and JSON formats for downstream modifications by the user.

### Hierarchical display of EC numbers and taxonomic affiliations

EsMeCaTa uses the OntoSunburst package to graphically display lists of predicted EC numbers (https://github.com/AuReMe/Ontosunburst). OntoSunburst is a Python package designed to visualize a set of concepts within an ontology. Applied to a given set of EC numbers, it displays the proportion of each EC class according to the four classification levels of the EC ontology. The EC ontology has been extracted from the Expasy databases (https://ftp.expasy.org/databases/enzyme/enzclass.txt, Release : 29-May-2024).

The list of predicted ECs is used as input to the OntoSunburst package, which extracts the EC ontology subgraph associated with that particular list. This subgraph is then plotted as a sunburst graph using the Plotly library [85]. The size of the sunburst patches corresponds to the proportion of the EC subclass in the list.

Similarly, the taxonomic sunburst provides a representation of the taxa diversity in the input taxonomic affiliation dataset. The proportion of each taxonomic group is represented with patch proportions. Each taxon selected by EsMeCaTa is coloured according to its taxonomic rank. Otherwise it appears in grey. The complete lineage of each taxon is retrieved from the NCBI taxonomy using the Python package ete3.

### Function enrichment analysis

The *esmecata_gseapy* command performs an enrichment analysis and automatically identifies the predicted functions that are enriched in a given taxon, using GSEApy [36] and Orsum [37]. Labels of annotations are retrieved from the Expasy ENZYME [86, 87] database for the EC numbers and from the Gene Ontology [88] database for the GO Terms. The enrichment analysis is performed by replacing the gene names by the taxa names given as input to EsMeCaTa. A pseudo GMT (Gene Matrix Transposed) file is then created with annotation IDs in the first column, annotation label in the second column and the list of observation name (the names of the taxa containing the annotation) in the following columns. The enrichr module of the GSEApy package uses the GMT file and the list of taxa of a phylum (by default) to find the annotations enriched in that phylum compared to the annotations present in the whole community. It creates one list of enriched terms per group. These lists of enriched annotations are finally provided to the Orsum method to extract a sublist of enriched annotations and compute several visualisation files.

## Comparison of EsMeCaTa predictions with MAGs and complete genomes

### Prediction assessment and statistical analysis

Confusion matrices were computed as follows for the different benchmarks considered. When comparing a feature predicted by EsMeCaTa (*i.e.*, an EC number or a consensus sequence) with a feature present in a genome (or, equivalently, in a MAG) considered as a reference, a true positive (TP) consisted of a feature found both in the reference genome and in the EsMeCaTa predictions. A feature that was present in the EsMeCaTa predictions but not in the reference genome was considered a false positive (FP). A feature missing from the EsMeCaTa prediction but present in the reference genome was considered as a false negative (FN). Then the performance metrics, precision, recall and F-measure, were computed as follow:

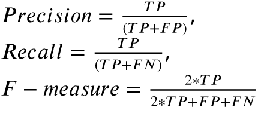

The measure distributions were visualised with boxplots and analysed with statistical tests. Due to the non-normality of the ANOVA residuals, Kruskal-Wallis tests were used in association with Dunn’s post-hoc tests (with Bonferroni correction for multiple tests). In Figure 3 B, for each taxonomic rank, Mann–Whitney U tests were performed (with Benjamini-Hochberg correction for multiple tests) to compare the performance of the two methods (EsMeCaTa against PICRUSt2).

To present the results of the post-hoc tests, compact letter displays were created using multcompView [89]. Each variable were assigned a letter that indicated if its mean was different from the ones of the other considered variables. If two variables shared the same letter, their mean were not statistically different whereas if they had different letters, their means were statistically different. Furthermore, compact letter display ranked the variables from the highest mean to the lowest mean.

Figures and statistical tests were computed using R version 4.4.1 [90] with the packages ggplot2 version 3.5.1 [91], FSA version 0.9.5 [92], rcompanion version 2.4.36 [93], multcompView version 0.1.10 [89] and tidyverse version 2.0.0 [94]. Linear model was made with stats package of R version 4.4.1 [90].

### Benchmarking the impact of 𝑅 cluster filtering threshold on EC prediction

Five runs of EsMeCaTa were performed on the *Ectocarpus sp. microbiota dataset* to test the effect of the representativeness threshold 𝑇_𝑟_. For each run a different value for 𝑇_𝑟_ was used: 𝑇_𝑟_ = 0, 𝑇_𝑟_ = 0.25, 𝑇_𝑟_ = 0.5, 𝑇_𝑟_ = 0.75 and 𝑇_𝑟_ = 0.95 (option *–threshold* of *esmecata clustering*). Annotations of the genome or the MAGs were predicted using eggNOG-mapper version 2.1.9 with eggNOG database version 5.0.2. The EC predictions from EsMeCaTa were then compared with the annotation of the genomes and MAGs in order to compute precision, recall and F-measure (see above).

### Assessing EC number and GO Term predictions compared to MGnify metagenomes

EsMeCaTa was applied to the *MGnify dataset* to evaluate its predictive performance on real environmental data. The taxonomic affiliations of the MAGs were used as input to EsMeCaTa. The annotations of the MAGs were retrieved from the corresponding eggNOG-mapper files in the MG- nify database. Then the EC numbers and the GO terms predicted by EsMeCaTa were compared with the EC and GO terms contained in the annotation file of the MAG (see above).

### Benchmarking EsMeCaTa and PICRUSt2 EC number predictions against MGnfiy metag- neomes

The predictions of EsMeCaTa were compared with the predictions of PICRUSt2 [8, 9] on the *MGnify dataset*. For this purpose, 16s rRNA sequences were extracted from the rRNA fasta files provided with each genome in the MGnify genome catalogs. If more than one 16S rRNA sequences was annotated in a genome, the longest one was selected as the representative. The PICRUSt2 (version 2.5.2) script "place_seqs.py" was then run to place the sequences within the PICRUSt2 database phylogenetic tree, creating a tree with the 16S rRNA sequences as new leaves. This tree was then passed to the PICRUSt2 script "hsp.py" to perform hidden-state prediction and predict the EC numbers. These EC numbers were compared with those in the annotation file of the MGnify MAGs, as described above in the "Measure of performance" section. Then the F-measures computed with predictions from PICRUSt2 were compared with the ones computed by EsMeCaTa in Figure 3 B.

### Evaluating the quality of consensus proteomes compared to MGnify metagenome proteins

A comparison was made between the consensus protein sequences predicted by EsMeCaTa and the protein sequences included in the MAGs. For each MAG, Diamond version 2.1.9.163 [51] was run on the MAG fasta file and on the consensus sequences predicted by EsMeCaTa. Two runs were performed, a first run with the MAG as query and the EsMeCaTa consensus proteome as reference and, reciprocally, a second run with the EsMeCaTa consensus proteome as query and the MAG as reference. The identified matches were filtered using an e-value greater than 1𝑒 − 05, a sequence identity greater than 40% and an alignment coverage greater than 50%, according to [41].

In order to test the similarity between the predicted consensus proteome and its MAG counterpart, two metrics were used, the Percentage of Conserved Proteins (POCP, [41]), and the Reciprocal Best Hits (RBH, [95]). First, the Percentage Of Conserved Proteins corresponds to the addition of the number of matches of the MAG to the EsMeCaTa proteome plus the number of matches of the EsMeCaTa proteome to the MAG, divided by the total number of sequences contained in both MAG and EsMeCaTa proteomes.

A confusion matrix was computed using RBH. An RBH was identified when, for two proteins (one from the MAG and one from the EsMeCaTa proteome), each protein matches the other as its best scoring match in the other proteome. An RBH was considered as a true positive (TP), a protein in the MAG without an RBH was considered as a false negative (FN) and a protein in the EsMeCaTa proteome without an RBH was considered as a false positive (FP). These measures are considered to compute precision, recall and F-measure (see above).

## Application of EsMeCaTa on a biogas reactor microbial communities

### Exploring specific functions of phyla from the community

The command *esmecata_gseapy* was used on the output of the run of *esmecata workflow* to identify functions enriched in a phyla compared to the all community. To this end, the EsMeCaTa version pre-release 0.5.0 was used with gseapy version 1.1.2. Orsum version 1.7.0 was used as to create Figure 4 B.

### Analysis of methanogenic pathways

An overview of several metabolic pathways of the methanogenesis was generated by combining searches in metabolic databases (MetaCyc version 28.0 [96, 97], KEGG version 110 [98, 99, 100], ENZYME Release of 29-May-2024 [86]) and literature [62, 101]. From MetaCyc, two pathways were used as reference: one consuming acetate (acetotrophic methanogenesis, MetaCyc Id: METH-ACETATE-PWY) and the other consuming H_2_ and CO_2_ (hydrogenotrophic methanogenesis, MetaCyc Id: METHANOGENESIS-PWY). METH-ACETATE-PWY was modified as some Archaea used *acs* instead of *pta* and *ackA* at the beginning of this pathway [59]. A pathway from methanethiol was also added to verify the predictions for *Candidatus Methanofastidiosum* [62].

### Homology search using the consensus protein sequences

Sequence homology searches against UniProt and KEGG for the methanogenic pathways and using proteins from literature for the cellulosome complex were carried out using the consensus proteomes predicted by EsMeCaTa. Diamond version 2.1.9.163 [51] was used for aligning the consen- sus proteomes from EsMeCaTa and the reference sequences. A first batch of reference sequences were made by mapping UniProt IDs and EC numbers from the ENZYME database flat files [86] (Release of 29-May-2024). A second set of reference sequences were retrieved from KEGG Orthologs (KO) [102, 103] associated with the EC number (KEGG Release 110.0). For each KO, a python script (using bioservices) extracted the protein sequences associated with the reference articles from its KEGG page. For the cellulosome complex, the dockerin and cohesin protein sequences from [50] were used as references. The identified matches were filtered using the ultra-sensitive option of Diamond, an e-value greater than 1𝑒 − 05, a sequence identity greater than 30% and an alignment coverage greater than 80%. Then, the resulting matches were filtered according to the RBH procedure. Two alignments were performed with Diamond, a first run with the reference protein sets as query and the EsMeCaTa consensus proteome as reference and, reciprocally, a second run with the EsMeCaTa consensus proteome as query and the reference protein sets as reference. Then a match between a protein from EsMeCaTa and a reference protein is kept only if each of the protein finds the other one as its best scoring match. Matches found with SwissProt were shown as stars and matches with KEGG Orthologs as circles (Fig 5).

### Impact of methanogenic OTU abundances on biogas production

To study the impact of trophic groups associated with methanogenic pathways on biogas production, the abundances of their OTUs were used. First, for each trophic group, the abundances of each OTU contained in them were summed. Then this summed abundance was normalised with a z-score normalisation across the different time points for each group. This normalised abundance was plotted. To decipher the cumulative effect of each group, a linear model was fitted from the non-normalised abundance of each group with R stat package version 4.4.1 [90].

## Additional Files

Sup File 1.

**Toy example dataset.** A tabulated file showing the thirteen taxa selected as toy example.

Sup File 2.

***Ectocarpus sp.* microbiota dataset.** A tabulated file indicating the 45 taxa from the *Ectocarpus sp*.

microbiota.

Sup File 3.

**MGnify dataset.** A tabulated file showing the the 3,664 taxa selected from MGnify.

Sup File 4.

**Methanogenic reactor dataset.** A tabulated file containing the 445 taxa sequenced from the experimental methanogenic reactor and their absolute abundances at the different time points of measure.

Sup File 5.

**Methanogenic reactor metabolite measures.** An excel file indicating the measures of the metabolites (biogas, fatty acids) in the methanogenic reactor.

Additional file 6.

**Additional pdf file on EsMeCaTa runs, toy example dataset analysis and methanogenic reactor experiments.** The additional file contains detailed description of EsMeCaTa runs (Sup Tables S1 and S2 for dependencies and runtimes on the experiments), further information on toy example dataset (Sup Fig S1), on validation dataset (Sup Fig S2) and on the methanogenic reactor experiments (Sup Fig S3-S10).

## Acknowledgments

We acknowledge the GenOuest bioinformatics core facility (https://www.genouest.org) for provid- ing the computing infrastructure. We are grateful to the INRAE MIGALE bioinformatics facility (MIGALE, INRAE, 2020. Migale bioinformatics Facility, doi: 10.15454/1.5572390655343293E12) for providing help and/or computing and/or storage resources.

## Author Contributions

Contributions were assigned according to the CRediT classification: Arnaud Belcour: Conceptualization, Data curation, Formal Analysis, Investigation, Methodology, Software, Supervision, Validation, Visualization, Writing - original draft, Writing - review and editing Pauline Hamon-Giraud: Methodology, Software, Visualization, Writing - original draft, Writing -review and editing

Alice Mataigne: Methodology, Software, Visualization, Writing - original draft, Writing - review and editing

Baptiste Ruiz: Software, Writing - review and editing Yann Le Cunff: Formal Analysis, Validation

Jeanne Got: Data curation, Writing - review and editing Lorraine Awhangbo: Investigation

Mégane Lebreton: Investigation

Clémence Frioux: Software, Writing - review and editing Simon Dittami: Validation, Writing - review and editing

Patrick Dabert: Conceptualization, Data curation, Formal Analysis, Funding acquisition, Investigation, Methodology, Project Administration, Resources, Visualization, Writing - original draft, Writ- ing - review and editing

Anne Siegel: Conceptualization, Supervision, Data curation, Formal Analysis, Funding acquisition, Investigation, Methodology, Project Administration, Software, Supervision, Validation, Visualization, Writing - original draft, Writing - review and editing

Samuel Blanquart: Conceptualization, Supervision, Data curation, Formal Analysis, Funding acquisition, Investigation, Methodology, Project Administration, Software, Supervision, Validation, Visualization, Writing - original draft, Writing - review and editing

## Data availability

The code of EsMeCaTa is available at: https://github.com/AuReMe/esmecata. Additional files for validation and visualisation are available in a

Zenodo archive: https://doi.org/10.5281/zenodo.14502342. Thesequencing data for the methanogenic reactor have been deposited in the European Nucleotide Archive (ENA) at

EMBL-EBI under accession number PRJEB83808.

