## Additional File 6 for "Estimating consensus proteomes and metabolic functions from taxonomic affiliations"

#### EsMeCaTa experiments

##### Runtimes on the different datasets

The time taken by the three steps of EsMeCaTa on the different datasets presented in the article is shown in Supplementary Table S1. All the experiments were performed with 10 cores and 60 Gb of RAM on a HPC.

The dataset *E. siliculosus microbiota* was splitted in two with data coming from the article by Burgunter et al. [4] and another one with data from KleinJan et al [13].

**Table S1.** Runtimes (in minutes) of EsMeCaTa steps for the different datasets with the number of taxonomic affiliations and R threshold used.

| Dataset | Number of taxonomic affiliations | R threshold | EsMeCaTa steps (in minutes) Proteomes | Clustering | Annotation |
|---|---|---|---|---|---|
| *Toy example* | 13 | 0.5 | 23.6 | 62.4 | 157 |
| Burgunter et al. | 46 | 0<br>0.25<br>0.5<br>0.75<br>0.95 | 62.7 | 97.3<br>47.9<br>47.9<br>53.8<br>43.9 | 1650.2<br>497.4<br>340<br>291.9<br>168.6 |
| Kleinjan et al. | 71 | 0<br>0.25<br>0.5<br>0.75<br>0.95 | 46.7 | 109.7<br>51.6<br>71.7<br>44<br>112.7 | 2017<br>397<br>340.4<br>513.7<br>479.9 |
| *Honeybee* | 627 | 0.5 | 74 | 39 | 638 |
| *Human Oral* | 1225 | 0.5 | 82.6 | 65.9 | 954.1 |
| *Marine* | 1504 | 0.5 | 218.9 | 210.5 | 2095.5 |
| *Pig Gut* | 3972 | 0.5 | 146.9 | 164.9 | 2436.1 |
| *Methanogenic reactor* | 445 | 0.5 | 107.5 | 95.3 | 953.6 |

#### Metadata and dependencies

EsMeCaTa runs on the *E. siliculosus microbiota*, *MGnify* and *toy example* datasets were performed in similar environments: Python 3.9.7, esmecata Pre-release 0.5.0, biopython 1.80, ete3 3.1.2, pandas 1.3.2, requests 2.26.0, SPARQLWrapper 1.8.5, mmseqs 14.7e284, eggnog-mapper 2.1.9 and eggnog database 5.0.2.

Run for the *methanogenic reactor* was performed with another environment: Python 3.12.1, esmecata Pre-release 0.5.0, biopython 1.83, ete3 3.1.3, pandas 2.2.0, requests 2.31.0, SPARQLWrapper 2.0.0, mmseqs 14.7e284, eggnog-mapper 2.1.9 and eggnog database 5.0.2.

The UniProt and NCBI Taxonomy database versions used in the article are shown in Supplementary Table S2.

**Table S2.** Version of database used for the datasets.

| dataset | UniProt | NCBI Taxonomy |
|---|---|---|
| *Toy example* | 2023_04 | 09-2023 |
| *E. siliculosus microbiota* | 2023_02 | 04-2023 |
| *Honeybee* | 2023_05 | 12-2023 |
| *Human Oral* | 2023_05 | 12-2023 |
| *Marine* | 2023_05 | 12-2023 |
| *Pig Gut* | 2023_05 | 12-2023 |
| *Methanogenic reactor* | 2024_01 | 01-2024 |

#### Visualisation of EsMeCaTa results for the toy example dataset

The distribution of EC numbers shown in Panel B of Figure 2 was further analysed. The distribution of classes was visualised in Sup Figure S1.

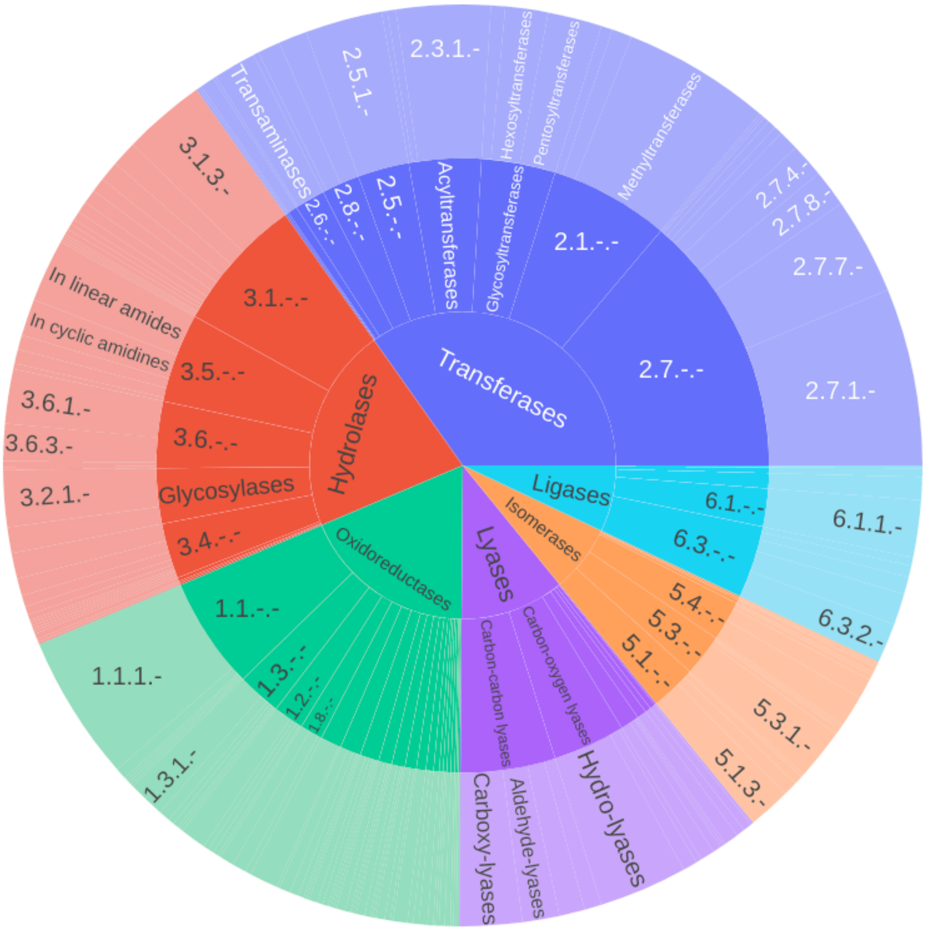


**Figure S1.** Distribution of the predicted functions (ECs) for genus *Escherichia*, the names corresponding to the first EC digits are indicated at the sunburst chart centre, internal and external rings respectively indicate the names or the numbers of second and third EC digits, and fourth EC digits are not shown. For the sake of readability, long EC names are replaced by the EC numbers, and ECs representing less than 1% of the total predicted EC number are not shown.

#### *MGnify dataset*: Discrepancies in prediction quality according to the datasets

Results presented in Figure 3 displays F-measures according to taxonomic ranks. Another impacting variable was the origin of the dataset as shown in Sup Figure S2. There was a significant difference in the distribution of F-measure according to the dataset. Datasets Honeybee and Human oral exhibited better F-measures than the Marine and Pig gut datasets. This could be explained by better sequencing technologies for some datasets and the fact that some environments has been more studied thus more genomes are available leading to better predictions.

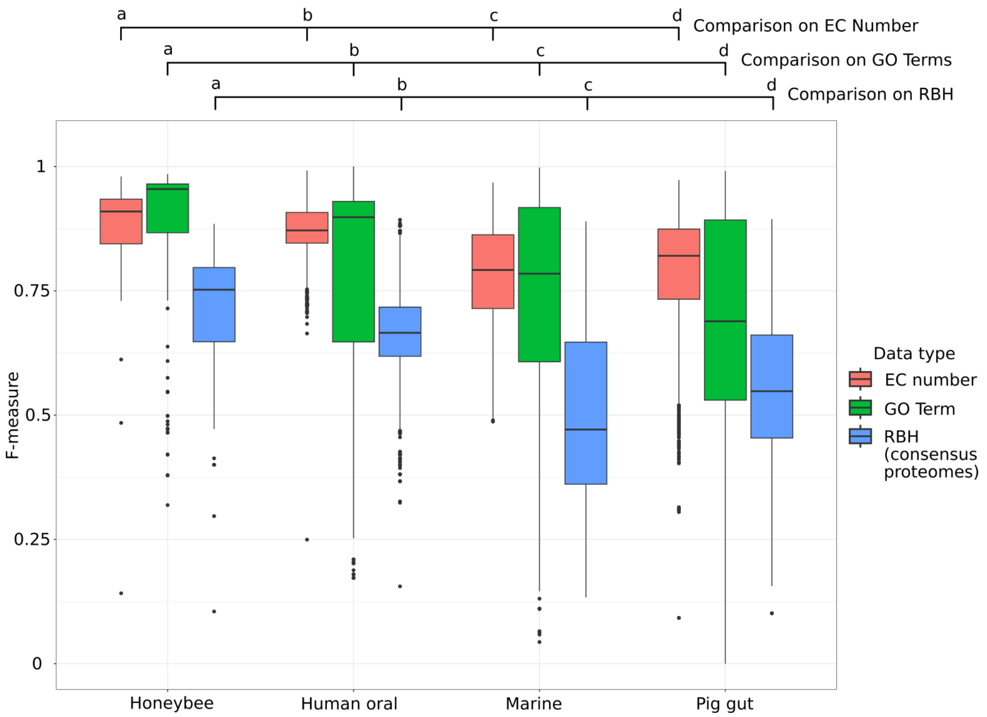


**Figure S2. Benchmark of EsMeCaTa predictions against 3,664 MAGS from MGnify database according to dataset.** F-measures computed from EC number, GO Terms and Reciprocal Best Hits (RBH) between EsMeCaTa proteomes and their associated MAGs (used as the reference).

### Experiments on a methanogenic reactor: operation and microbial community characterization

#### Materials and Methods of methanogenic reactor experiment

The aim of this experiment was to analyze changes in the microbial community of a laboratory pilot reactor in response to major substrate changes.

Process operation was published in [1]. A 35L working volume anaerobic continuously stirred tank reactor (CSTR) was run for 195 days under mesophilic conditions (38°C) in four successive experiments designed to study different feeding stresses for anaerobic digestion (Figure S3). All substrates were prepared every week, ground and stored at 4°C to feed the reactor once a day. The pH and biogas production of the process were monitored online. Biogas composition was analyzed once a day by gas chromatography. Digestate composition (organic matter, nitrogen, volatile and long chain fatty acids) was determined once a week. Every week, 15 mL of digestate were collected from the reactor, centrifuged at 12,000 g for 10 min to recover the pellet that was frozen at -20°C.

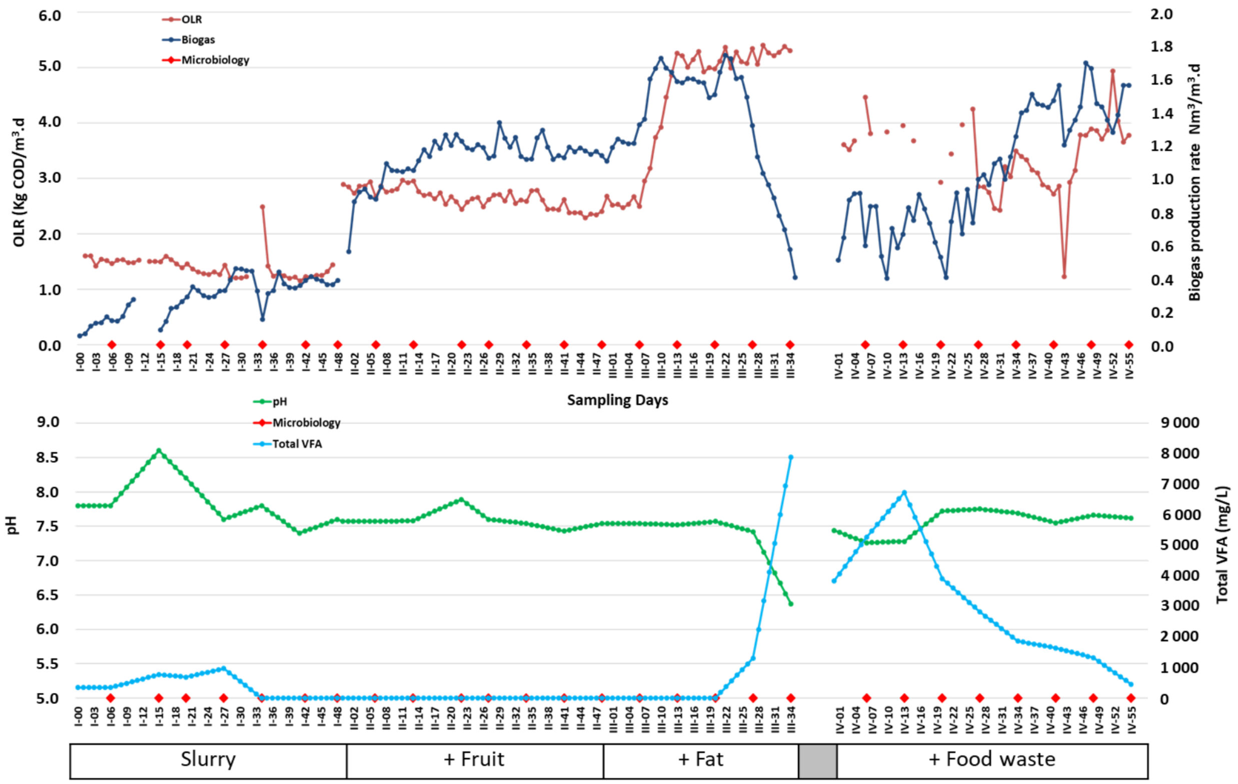


**Figure S3.** Top, organic loading rate and biogas production of the methanogenic reactor during the 195 days of the experiment and the 4 different feeding periods: I-00 to I-48 for slurry, II-00 to II-48 for slurry + fruits, III-00 to III-34 for slurry + fat and IV-00 to IV-55 for slurry + food waste. Period in grey means 7 days without feeding. Bottom, pH and VFA concentration. Red diamonds are the days of sampling for microbial community metabarcoding.

Total genomic DNA was extracted from about 250 mg of frozen pellet using the NucleoSpin® Soil DNA extraction kit (Macherey-Nagel, USA) as recommended by the manufacturer. The microbial community characterization was performed by Ion Torrent high throughput DNA sequencing at the metagenomic platform of the UR1461 PROSE of INRAE (Antony, France). The archaeal and bacterial hypervariable V4-V5 regions of the 16S rRNA genes were amplified by PCR using the so-called universal primers 515F (5'- Ion A adapter-Barcode-GTGYCAGCMGCCGCGGTA-3') and 928R (5'-Ion trP1 adapter-

CCCCGYCAATTCMTTTRAGT-3') as described in [15] The resulting amplicons were purified, quantified and sequenced at the metagenomic platform of the UR1461 PROSE of INRAE (Antony, France) according to manufacturer's instructions. Sequencing was performed on an Ion Torrent Personal Genome Machine using Ion 316 Chip V2 (Life Technologies) and Ion PGM Hi-Q View Sequencing Kit (Life Technologies). The sequencing reads were processed with the FROGS pipeline [9] following the authors' recommendations on the MIGALE Galaxy instance (INRAE, Jouy-en-Josas, France). Taxonomic affiliation of the OTUs was done on 16S SILVA Pintail100 database [18].

#### Results of methanogenic reactor experiment

Figures S3 and S4 show respectively the principal parameters of process operation and the relative abundance of the major microbial families during this period. Only the families making up to 0.1% of the complete microbial community are presented. The microbial community was dominated by members of the *Bacteroidota*, *Bacillota*, *Cloacimonadota*, *Spirochaetota* and *Chloroflexi* phyla for the bacterial domain. While the archaeal sequences belonged to methanogenic genera of the *Euryarchaeota* and *Halobacterota* phyla. This microbial community structure is classical from anaerobic digestion processes [5, 14, 21].

##### Process stabilization

The reactor was first stabilized at a low organic loading rate (OLR) of 1.4 +/- 0.2 Kg COD.m-3.d-1 with a feed based on pig slurry completed with horse feed for 48 days (Fig. S3, Slurry, Exp. I-00 to I-48). This feed provided the inoculum and nutrients required to ensure good anaerobic digestion and a biogas production of about 0.37 +/- 0.07 Nm3/m3.d-1.

This first phase of reactor stabilisation did not strongly impacted the bacterial community that remained dominated by the *Dysgonomonadaceae* and the *Sphingobacteriales* ST-12K33 at about respectively 15% and 10% relative abundance. The former family contains fermentative species able to degrade complex carbohydrate and proteinaceous substrates. The *Sphingobacteriales* activities are poorly documented but they have been observed in several anaerobic digestion processes treating food waste. The other next dominant families were the *Anaerolineaceae* (species *Brevefilum fermentans*), the *Prolixibacteraceae* and DTU014uk whose relative abundance decreased from about 8% to 2-3% during this phase. The *Anaerolineaceae* degrade sugars and proteins while the *Prolixibacteraceae*

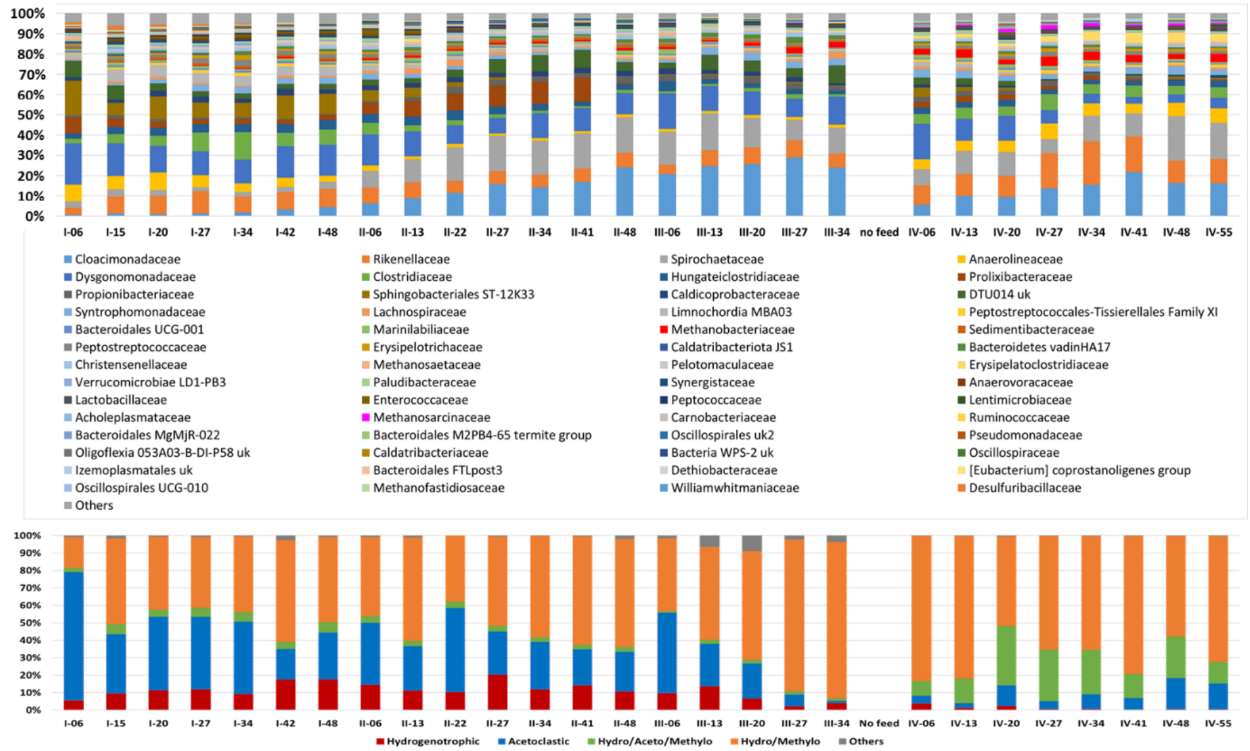


**Figure S4.** Top, relative abundance of the bacterial and archaeal families that are present at more than 0.1% of the complete microbial community. Bottom, relative abundance of the archaeal families clustered according to their putative metabolic pathways of methanogenesis in the literature, with Hydro, aceto and methylo meaning respectively hydrogenotrophic, acetotrophic and methylotrophic.

---

uses alcohols, VFA and sugars as carbon sources. The DTU014uk uncultured microbial group is also poorly documented but is suspected to perform the synthrophic oxidation of acetate into CO2 and H2. On the opposite, the *Clostridiaceae* (*Clostridium sensu stricto 1*) and *Rikenellaceae* (RC9 gut group) relative abundance increased from 2-3% to 8-9%. Both families are involved in the anaerobic degradation of complex carbohydrates and proteins to produce VFA and alcohols. The *Clostridium sensu stricto 1* species possesses the Wood-Ljungdahl pathway and can convert CO2 into acetate. During all this period, the methanogenic archaea remained below 2.2% abundance of the total microbial community. However, focusing on this domain shows a shift from a domination by the acetotrophic *Methanosaeta* representing about 70% of the archaea on day I-06 to a community a share at about 50% with the hydrogenotrophic and methylotrophic *Methanobacterium*. This apparent shift from an acetotrophic to hydrogenotrophic methanogenesis is accentuated by the increased relative abundance of the hydrogenotrophic *Methanocorpusculum* from 0.04% to 0.14% relative abundance. Overall, this microbial community structure suggests the existence of all the known metabolic pathways needed for organic matter anaerobic digestion with a last step involving firstly acetotrophic methanogenesis but with a possibly increasing share of hydrogenotrophic methanogenesis.

#### Addition of easily degradable carbohydrates and performance increase

At this stage, model substrates were added to increase the OLR. Addition of ground apples to the feed, which is an easily digestible carbohydrate substrate, increased the OLR to 2.6 +/- 0.2 kg COD.m-3.d-1 and allowed to increase biogas production up to a steady state around 1.18 +/- 0.06 Nm3/m3.d-1 (Fig. S3, Fruit, Exp II-00 to II-48). This promoted the strong increase in the relative abundance of three bacterial families: the *Cloacimonadaceae* from 4% to 24%, the *Spirochaetaceae* (*Sphaerochaeta*) from 4% to 17% and to a lesser extent the comeback of *Prolixibacteraceae* and DTU014uk from 2-3% to 9-12%. This was on the detriment of the *Clostridiaceae*, the *Sphingobacteriales* ST-12K33 that almost disappeared from the dominant families. These bacterial shifts appear very logical since the addition of apples increased the share of easily degradable sugars in the feed of the reactor. This favoured the *Spirochaetaceae* that are known to ferment many sugars into acetate, formate and ethanol and the *Prolixibacteraceae* that uses alcohols, VFA and sugars as carbon sources. The *Cloacimonadaceae* also ferment many sugars, proteins and short chain fatty acids but this microbial group is also involved in the syntrophic oxidation of propionate into acetate and H2 and CO2, as long as there is H2 scavenger microorganisms, usually hydrogenotrophic archaea. In the archaeal domain, which represented between 1.25% and 2.48% of the microbial community, some fluctuations were observed but the acetotrophic methanogens remained subdominant at about 30% of the total archaea.

#### Addition of fat and process overloading

On the opposite, the addition of butter simulating the addition of fat from agri-food, increased the OLR rapidly up to 5.2 +/- 0.2 kg COD.m-3.d-1 leading first to a sharp increase of biogas production up to a steady state around 1.61 +/- 0.07 Nm3/m3.d-1. However, this OLR was too high to be managed by the process, resulting in a desynchronization between the fast growing fermentative bacteria that produce VFA and the slow growing methanogenic archaea that consume it. VFA started to accumulate in the reactor, reaching up to 8 g.L-1 of VFA in the digestate, inducing a pH drop to 6.4 and a biogas production failure (Fig. S3, Fat, Exp. III-00 to III-34). The feeding was thus stopped for 7 days.

Surprisingly, the addition of butter to the feed barely impacted the relative abundance of the dominant bacterial groups that remained relatively stables except for the *Prolixibacteraceae* that completely disappeared. The archaea however showed two behaviour. First, the proportion of archaea in the total microbial community increased up to 3.8% on day III-27, while the hydrogenotrophic and methylotrophic *Methanobacterium* represented about 81% of the total archaea.

#### Process recovery with slurry and food waste

Finally, the biogas production was recovered by reducing the OLR to 3.8 +/- 0.4 Kg COD.m-3.d-1 and feeding the reactor with a mixture of pig slurry, horse feed and food waste from a restaurant (Fig. S3 Food waste, Exp. IV-00 to IV-55). During this period, the reactor consumed the VFA and the biogas production stabilized around 1.43 +/- 0.12 Nm3/m3.d-1.

The bacterial community seems to stabilize with three dominant families whose relative abundance fluctuate between 10% and 20% but makes about 50% of the total microbial community: the *Spirochaetaceae* (*Sphaerochaeta*), the *Cloacimonadaceae* and the *Rikenellaceae*. The mean proportion of archaeal sequences in the total microbial community is about 5.3% with a peak at 7.1% on day IV-27. The hydrogenotrophic and methylotrophic *Methanobacterium* remains the dominant archaea representing 50%-70% of the total archaea but this process operation stage is marked by the increase in the versatile hydrogenotrophic, acetotrophic and methylotrophic *Methanosarcina* (8.6% to 33.9%) and the comeback of the acetotrophic *Methanosaeta* (2.6% to 18%).

### **Results of EsMeCaTa on the biogas reactor**

The global distribution of the EC numbers predicted at the community scale was examined with further details by extracting them from the function table. The distribution of the predicted functions over the taxa corresponded to an estimate of the environment EC numbers (Fig S6A) and GO terms (Fig S6B).

Some predicted functions were shared by all the taxa in the community, forming the core predicted functions in the environment. In contrast, most of the predicted functions were specific to one or few of the taxa (Fig S6A and B), and some might represent key functions in the environment. Regarding the example, some of these taxa specific predicted functions were linked to methanogenesis in archaea.

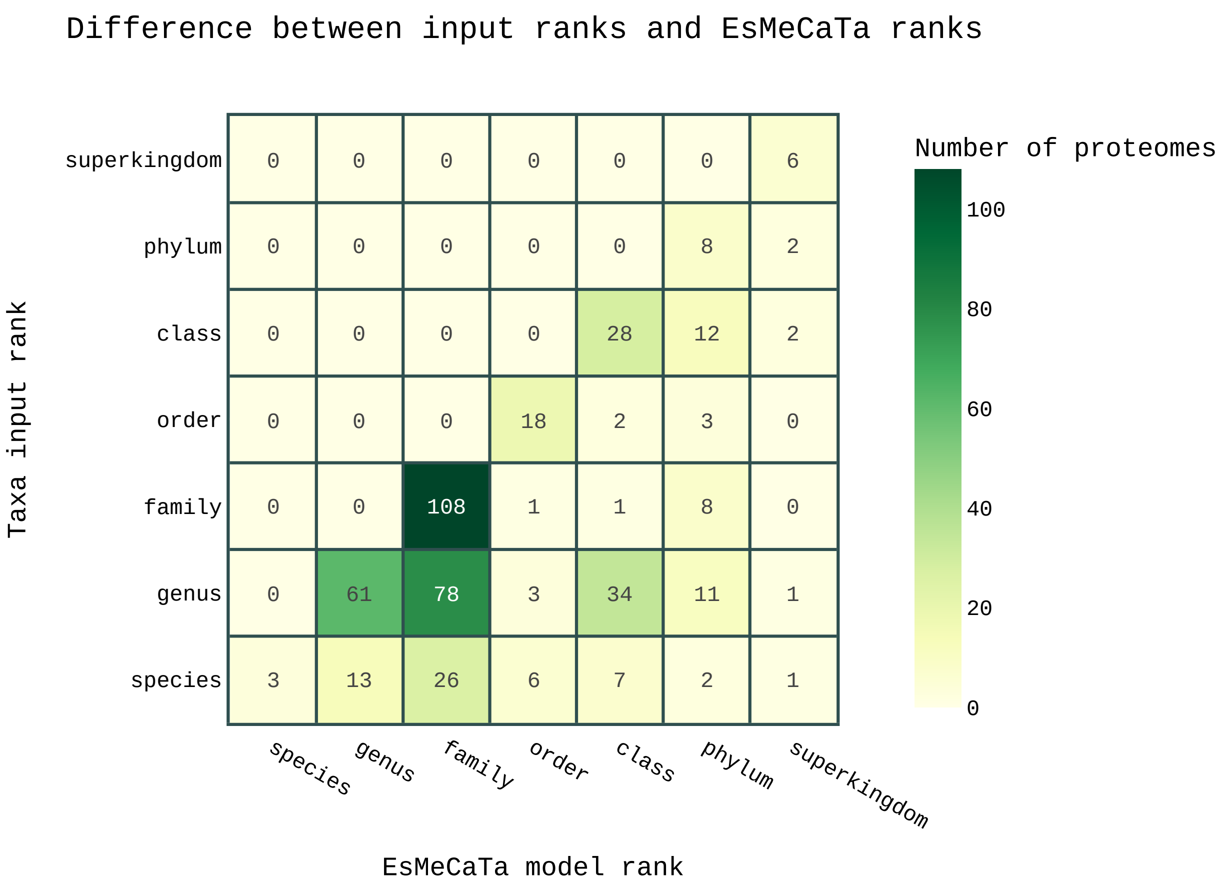


**Figure S5. A methanogenic reactor taxonomic diversity.** (A) Taxonomic diversity of 445 OTUs estimated from 16S rRNA amplicon sequencing of an experimental methanogenic reactor. (B) Number of OTUs with given input taxonomic ranks (table lines) and given ranks selected (table columns) according to the availability of proteomes in the UniProt proteome database. Almost all the OTUs have input ranks genus and higher, and the majority of the predictions are drawn for ranks family and higher, suggesting an organism diversity poorly characterised both at the 16S rRNA amplicon and at the proteome levels.

#### Detecting key functions of taxonomic groups of the methanogenic community

Specifically, an enrichment analysis using GSEApy [10] was performed (see Methods) to identify the enriched functions in each phylum present in the methanogenic reactor samples, compared to the functions in the whole community. Then the results were processed by Orsum [17] to filter the resulting list of enriched terms (see Methods). This yields a condensed view of enriched functions in the phyla present in the studied community (Fig S7). The taxa from the *Euryarchaeota* phylum present in the samples were all methanogenic Archaea (Fig S3 A). Consistently, two of the four enriched func-

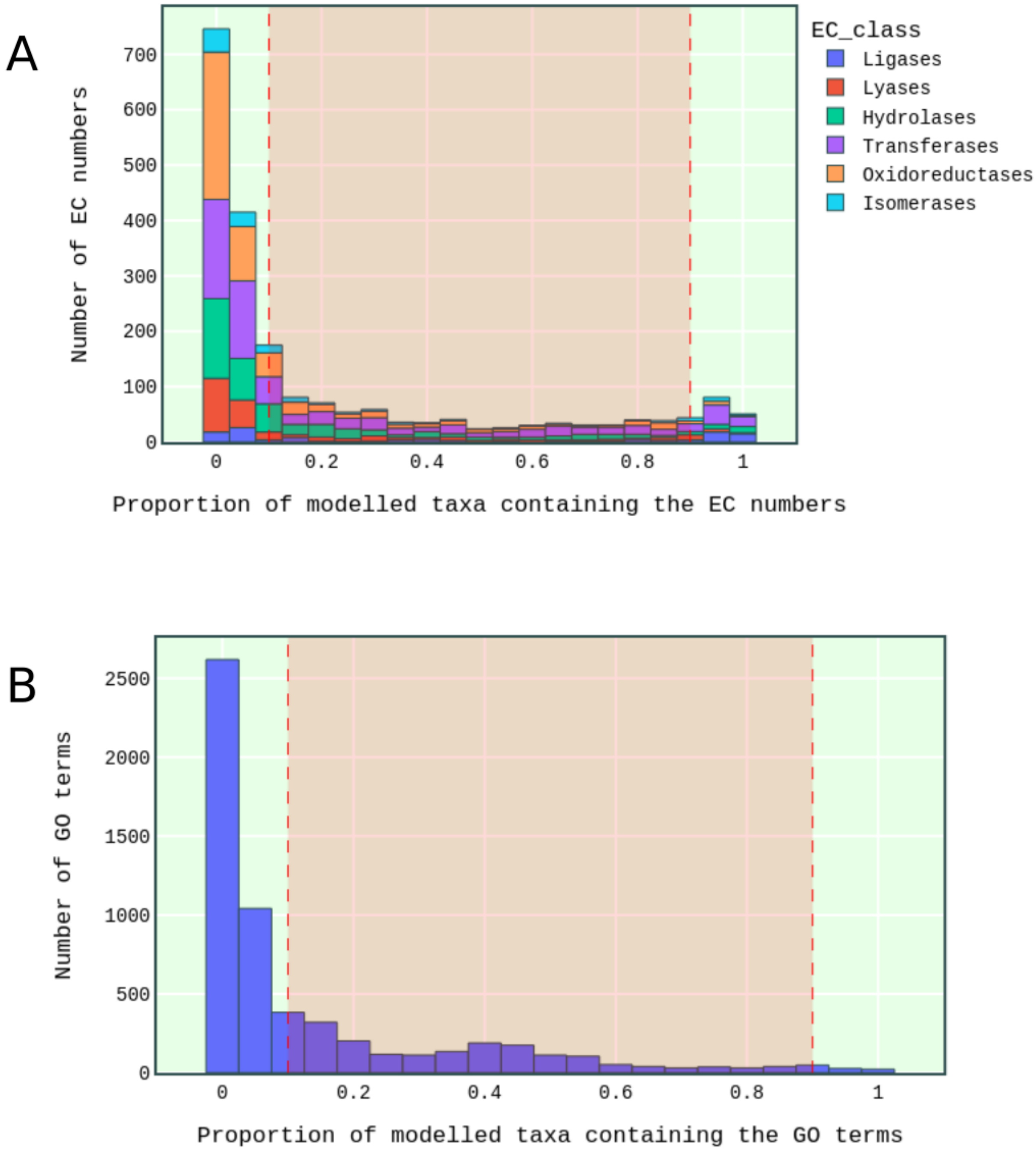


**Figure S6. Annotations predicted on the biogas reactor community dataset.** (A) Number of the predicted ECs according to the proportion of the taxa in the community sharing them. The number of main EC classes are represented with coloured stacks, the most widespread functions predicted in all the taxa are shown at the right bin, and specific functions predicted in one or a few taxa shown at the left bins. (B) Predicted GO terms shared across taxa.

tions associated with the *Euryarchaeota* were associated with methanogenesis, the EC number 2.8.4.1 catalysing the last step of methanogenesis and the EC number 2.1.1.245, a key enzyme in methanogenesis from acetate (Fig S7).

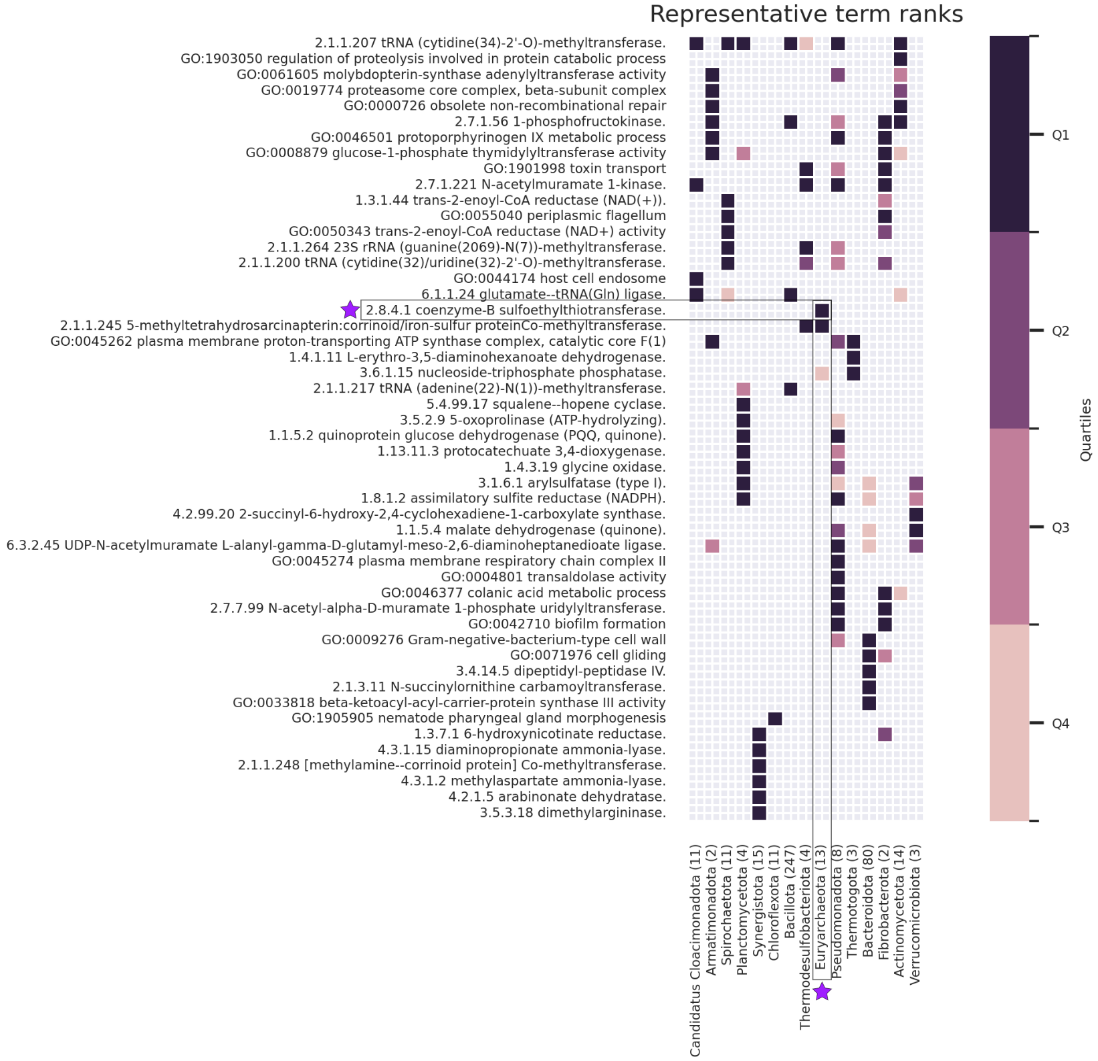


**Figure S7. Annotations (GO terms and EC numbers) enriched in the methanogenic reactor major phyla, compared to the whole community annotations.** The figure is produced using Orsum [17]: the enriched functions (GO Terms and EC numbers) are shown on the left figure side, coloured squares in the middle matrix indicate the taxa owning the enriched functions, and the colour scale on the right figure side indicates the enriched function rank, estimated with GSEApy[10] and binned into quartiles. Number of OTUs contained in the phyla are indicated in parenthesis after the phylum name. Purple star indicates the focus on the *Euryarchaeota* phylum and the EC number 2.8.4.1 associated with final step of methanogenesis. The most enriched annotations correspond to the most specific functions in a phylum compared to the functions in the whole community.

#### Detection of the cellulosome complex using EsMeCaTa consensus proteomes

The cellulosome catalyses the degradation of the cellulose and it is expected to be expressed in the methanogenic community. This complex system containing tens of enzymes [11] was not suitable described in metaoblic databases. For example, the reaction describing the degradation of cellulose to cellotetraose in MetaCyc (reaction RXN-14887) was modelled by as a sequence of reactions which were not associated with EC number or genes, therefore not predictable from genome analyses.

The consensus sequences of the homologous protein clusters provided by EsMeCaTa can be used to align to protein sequences of known functions using sequence homology search. This had been applied to the cellulosome to search for enzymes forming the structure of the complex (dockerin and cohesin). The dockerin and cohesin protein sequences from [2] were compared to the cluster consensus sequences estimated by EsMeCaTa using Diamond [3], showing that eight taxa in this dataset had predicted consensus proteomes containing related sequences (Fig S8 A, B). Genera *Acetivibrio* and *Ruminiclostridium* were predicted to possess both the dockerin and cohesin functions (Fig S8 C), in agreement with previous works on those taxa cellulosomes [7, 19].

#### Focus on methanogenesis and acetogenesis pathways

Following the analysis presented in Figure 5, other metabolic pathways were explored. Two pathways producing methane were considered (Fig S9 A): one consuming acetate (aceticlastic methanogenesis, MetaCyc Id: METH-ACETATE-PWY) and another consuming $H_2$ and $CO_2$ (hydrogenotrophic methanogenesis, MetaCyc Id: METHANOGENESIS-PWY). EC numbers from these pathways were retrieved from MetaCyc, manually curated and were queried into the EsMeCaTa raw predictions to reveal which taxa are predicted to achieve those functions involved in the pathways (see Methods). Another focus was made on part of the Syntrophic Acetate Oxidation (SAO, MetaCyc Id PWY-8480) pathway transforming acetate into $CO_2$, and part of the Wood-Ljungdahl pathway (WLP, MetaCyc Id CODH-PWY) reciprocally transforming $CO_2$ into acetate (Fig S9 A). The examined part of the WLP and SAO pathways involved the same four first reversible reactions as in aceticlastic methanogenesis [8] (Fig S9 B).

Concerning aceticlastic methanogenesis, the two starting EC numbers degrading acetate were predicted to be present in a broad range of bacterial taxa and in the methanogenic archaea genus *Methanosarcina* (Fig S10 A). This was consistent with the widely used reversible transformation of acetate into Acetyl-

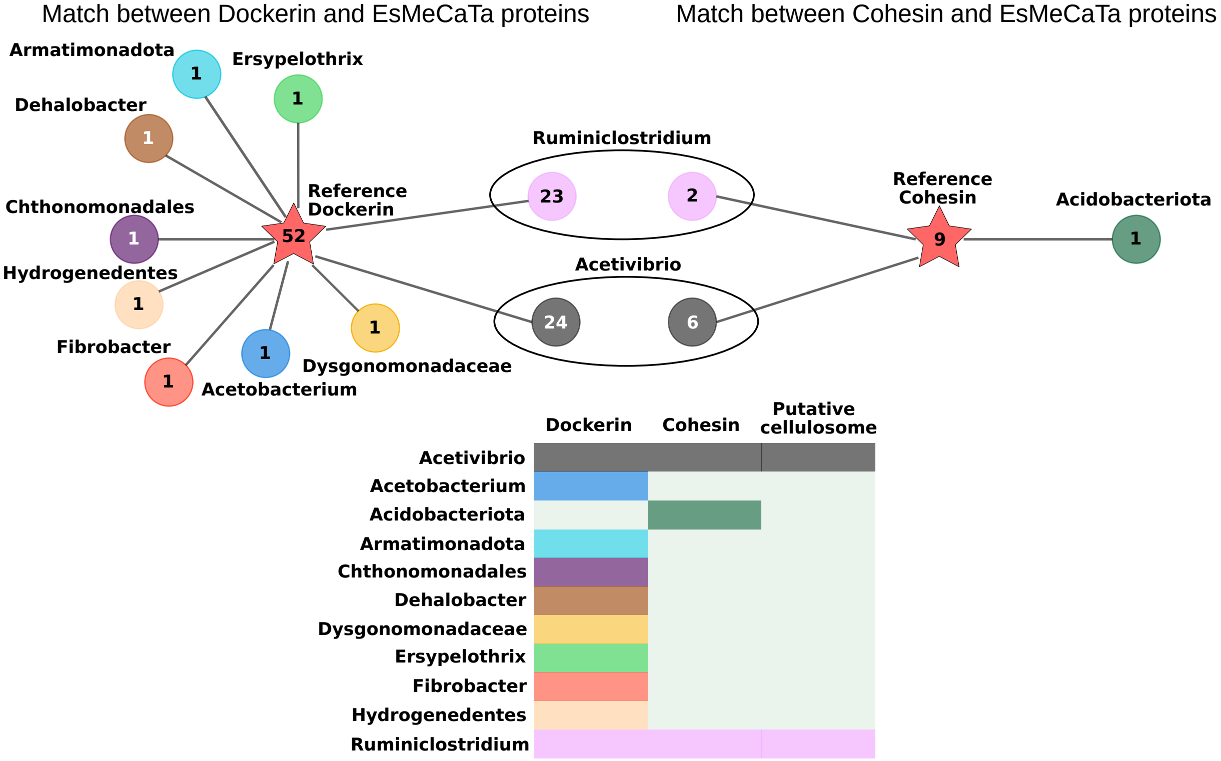


**Figure S8. Taxa predicted to contain dockerin and cohesin protein orthologs.** A set of dockerin and cohesin protein sequences was obtained from [2] and aligned with Diamond [3] for sequence homology with the cluster consensus sequences estimated by EsMeCaTa(red stars). Results were filtered according to RBH procedures. Number of sequences matching dockerin and cohesin respectively, in each taxon. Nodes correspond to the matching taxa, with their names on the right, and edges indicate the matches with the reference protein sequences. Taxa having both matches with dockerin and cohesin protein sequences are predicted to possess cellulosomes [2].

CoA in acetotrophic or acetogenic bacteria. Moreover this compound is involved into several pathways over than aceticlastic methanogenesis, such as SAO or WLP. In contrast only the final EC number of methanogenesis was found specifically in methanogenic archaea taxa (Fig S10 A), as expected in biogas reactor [5, 8]. Among them, the genus *Methanosarcina* showed the highest pathway completion, with one EC missing (EC: 2.3.1.169) over seven ECs considered (Fig S10 A). These organisms were consistently known to perform aceticlastic methanogenesis [6]. Moreover, among the sampled archaea *Methanothrix* (formerly *Methanosaeta*) is known to use acetate to perform methanogenesis [8]. Unlike in the *Methanosarcina* genus, it uses the reaction Acetyl-CoA synthase (EC: 6.2.1.1) to transform acetate into acetyl-CoA [12]. Congruently the two first reactions in aceticlastic methanogenesis were not predicted as present in the *Methanothrix* genus (Fig S10 A).

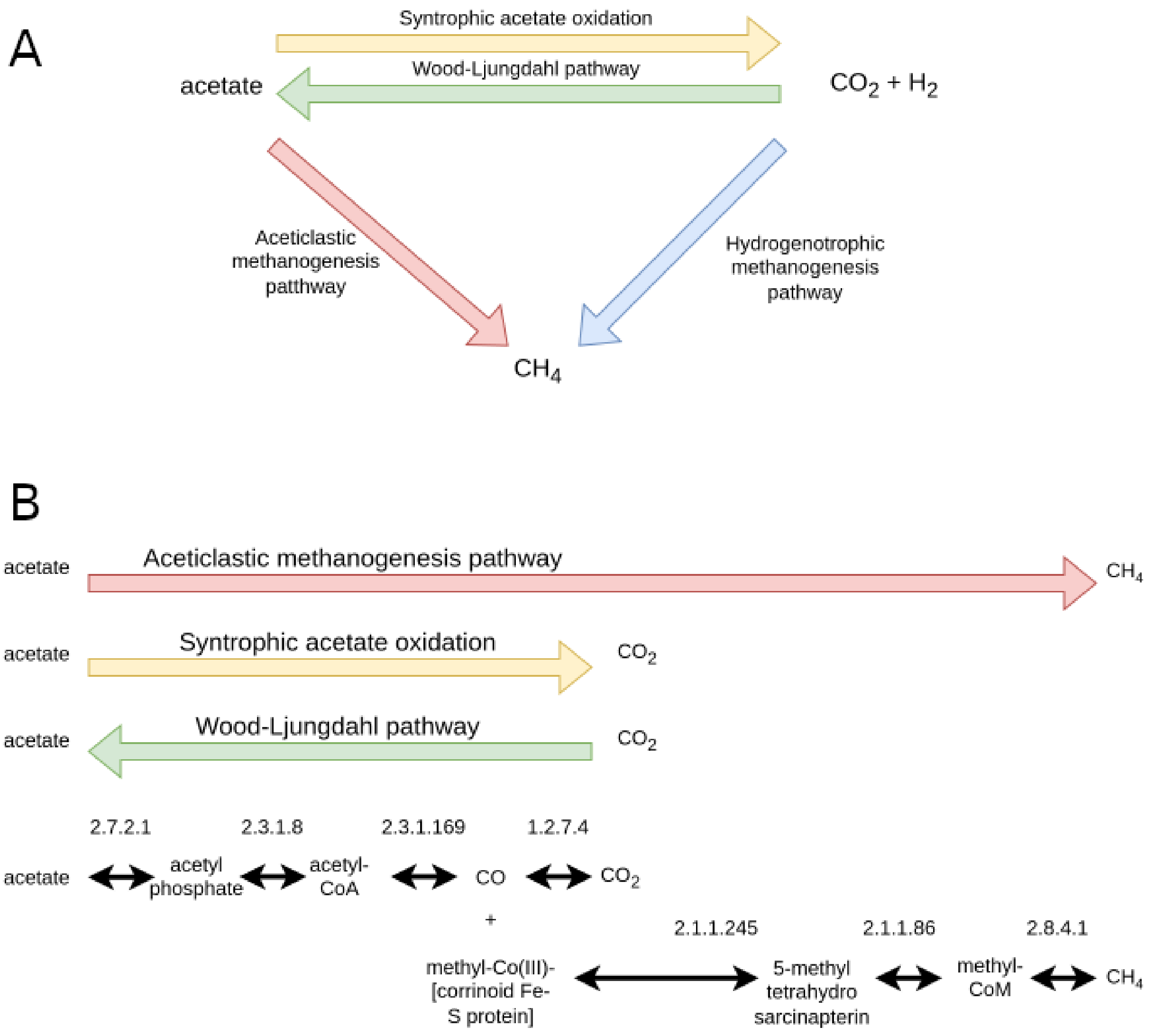


**Figure S9. Pathways present in methanogenic communities.** (A) Simplified of pathways producing acetate, or $H_2$ and $CO_2$, or $CH_4$. (B) Simplified representation of EC numbers involved in aceticlastic methanogenesis, syntrophic acetate oxidation (SAO) and Wood-Ljungdahl pathways (WLP). EC numbers and the key intermediate products were ordered as in the aceticlastic methanogenesis pathway described in *Methanosarcina*. Aceticlastic and hydrogenotrophic methanogenesis pathways are shown in red and blue respectively, and SAO and WLP pathways in yellow and green.

SAO is coupled with hydrogenotrophic methanogenesis in methanogenic environments [6, 8]. It is achieved by bacterial species from several classes [16]. In the class *Dethiobacteria*, the genus *Dethiobacter* is known to perform SAO [8]. EsMeCaTa predicted two of the four examined SAO reactions in this taxon (Fig S10 A, highlighted in yellow). Other taxa sequenced from the biogas reactor were known to perform the reverse process WLP. Reactions of the WLP have been shown to be present in the family

*Peptococcaceae* [5] and in the genus *Acetobacterium* [20]. In agreement with these studies EsMeCaT-apredicted the WLP EC numbers in these two taxa (Fig S10 A, taxa highlighted in green). Thus the EC numbers successively occurring during aceticlastic methanogenesis in archaea species could be partly predicted in some bacterial clades where they could be involved in other pathways, such as SAO and WLP. EC numbers from the hydrogenotrophic methanogenesis pathway (Fig S9 A, blue arrow) were all predicted in the archaea identified in the reactor community (Fig S10 B), except for *Methanofastidiosum*.

However, further refinements were required to complete pathways, such as concerning EC 2.3.1.169 mentioned in MetaCyc to apply in the aceticlastic methanogenesis, but which is not predicted by our method in genus *Methanosarcina* (Fig S10 A). Furthermore, the reaction 1.12.98.2 indicated in MetaCyc to apply in the hydrogenotrophic methanogenesis pathway was not predicted by our method in the methanogenic archaea. In the presented manually curated pathway and according to KEGG, it was replaced by the equivalent F420 dependant reaction, with EC 1.5.98.1 (Fig S10 B).

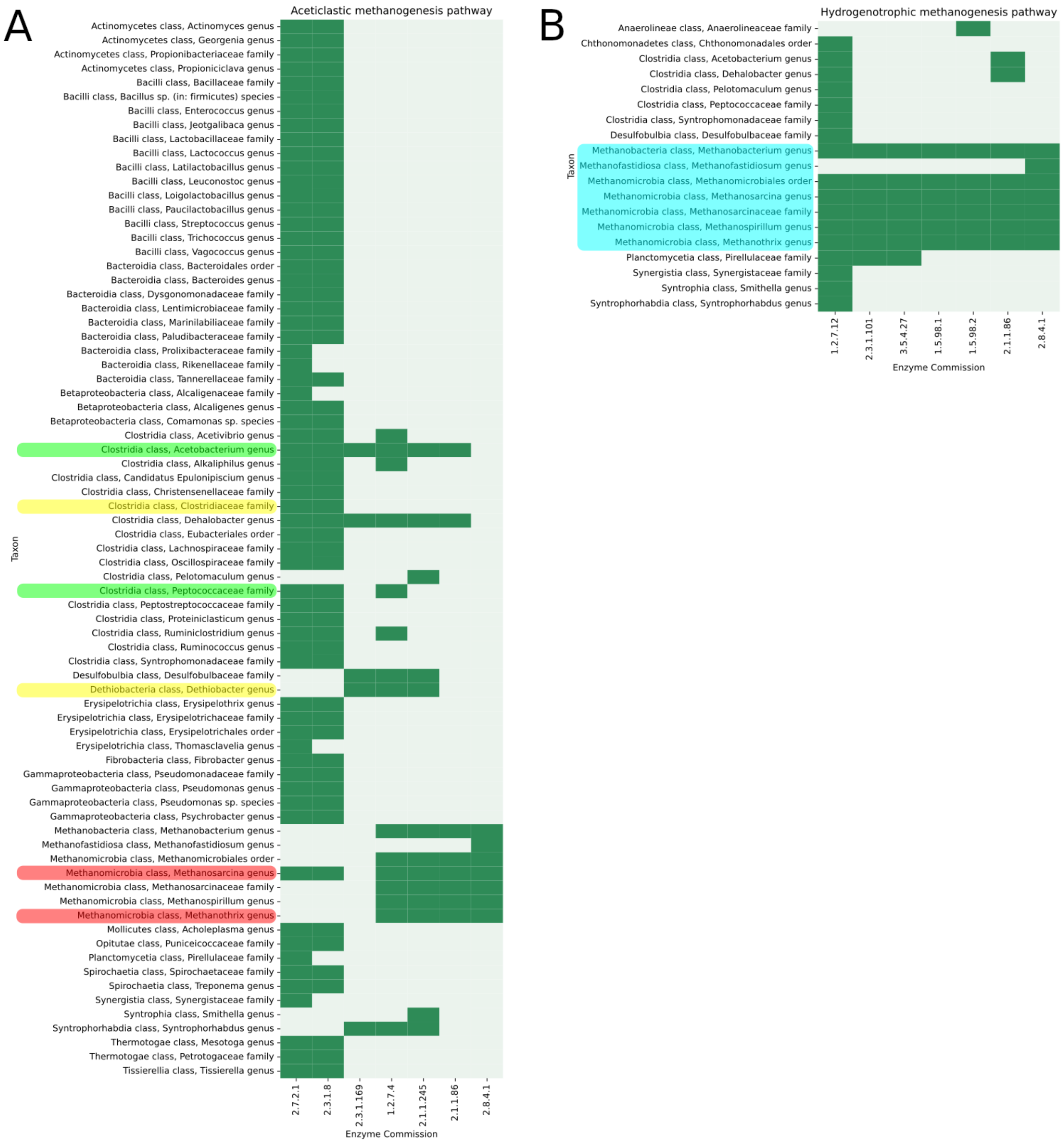


**Figure S10. Predicted EC numbers belonging to pathways involved in methanogenic communities.** (A) Taxa with predicted reactions from the aceticlastic methanogenesis pathway. (B) Taxa with predicted reactions from the hydrogenotrophic methanogenesis pathway. The ECs predicted in the taxa are shown with dark green cells, acetotrophic methanogenic archaea are highlight in red, hydrogenotrophic methanogenic archaea in blue, SAO performing bacteria in yellow, and WLP performing bacteria in green. The taxonomic class names are indicated for each taxa, followed by the taxa names and ranks. Predictions for phyla and higher taxonomic ranks are not shown.
